# A bacterial expression cloning screen reveals tardigrade single-stranded DNA-binding proteins as potent desicco-protectants

**DOI:** 10.1101/2023.08.21.554171

**Authors:** Jonathan D. Hibshman, Courtney M. Clark-Hachtel, Kerry S. Bloom, Bob Goldstein

## Abstract

Tardigrades are known for their ability to survive remarkable extremes, but relatively little is known about the molecular underpinnings that allow for their extraordinary stress tolerance. We devised an expression cloning approach to screen for novel protectants from desiccation-tolerant tardigrades. We expressed cDNA libraries from the tardigrades *Hypsibius exemplaris* and *Ramazzottius varieornatus* in *E. coli* and subjected the bacteria to desiccation. Sequencing the populations of surviving bacteria revealed enrichment of mitochondrial single-stranded DNA-binding proteins (mtSSBs) from both tardigrade species. Expression of mtSSBs in bacteria improved desiccation survival to similar levels as some of the best protectants known to date. The DNA-binding activity of the oligonucleotide/oligosaccharide binding (OB) fold domain of mtSSB was necessary and sufficient to improve desiccation tolerance. By comparing the efficacy of mtSSBs from multiple organisms, we provide evidence that desicco-protection is a conserved feature of mtSSBs. More broadly, any OB-fold containing protein may act similarly to promote desiccation tolerance, likely by coating ssDNA to limit damage during desiccation. These results identify single-stranded DNA-binding activity as a potent contributor to extreme desiccation tolerance.

## Introduction

In 1702, Antonie van Leeuwenhoek described reviving dried ‘animalcules’ from desiccated sediment^1^. Researchers have since been fascinated with the phenomenon of anhydrobiosis – life without water. Desiccation causes extensive damage to cells, yet some animals, like those observed by van Leeuwenhoek, are able to survive extreme desiccation^2, 3^. Desiccation may be a selective pressure that can drive adaptations to other extremes, like the tremendous radiation tolerance of the bacterium *Deinococcus radiodurans*^4^. Desiccation tolerance has also been linked to pathogenicity of clinical isolates of bacteria^5^. Beyond understanding the basic biology of anhydrobiosis, the identification of effective protectants may prove useful for stabilizing biological materials, like medicines, cells, or tissues^6–8^. Still, the protectants and molecular mechanisms that facilitate desiccation tolerance remain largely unknown.

Tardigrades, also known as water bears, are recognized for their ability to survive extreme conditions including desiccation, high levels of radiation, and the vacuum of space^9–11^. The capacity of tardigrades to survive what most animals cannot suggests that they must harbor uniquely powerful protectants. There is great interest in discovering such molecules. Some approaches have effectively identified tardigrade protectants by isolating heat-soluble proteins that are particularly stable^12^, identifying DNA-binding proteins that can protect against DNA damage^13^, and using the transcriptional response of tardigrades to prioritize candidates^14^. To be useful for applications in biomedicine, a protectant must have potency alone in diverse, non-native contexts. We were encouraged by the examples of tardigrade cytosolic abundant heat soluble (CAHS) proteins that were sufficient to improve desiccation survival in bacteria and yeast and the damage suppressor (Dsup) protein which could improve radiation tolerance of human cells^13, 14^. We sought an unbiased screening method for new protectants that would not rely on criteria like transcriptional regulation during desiccation and would prioritize candidates with potentially broad applicability beyond tardigrades.

Here, we report the implementation of a bacterial expression cloning screen to identify proteins from tardigrades that can serve as desicco-protectants in *E. coli*. Our screens revealed that mitochondrial single-stranded DNA-binding proteins (mtSSBs) from two species of tardigrades are potent desicco-protectants. The protective ability of mtSSBs relies on their binding to DNA and is conserved across species. We propose a model in which coating ssDNA with protein can limit DNA damage to promote desiccation tolerance.

## Results

### Developing bacterial expression cloning as a functional screening approach to identify new protectants

To identify new desicco-protectants from tardigrades, we developed a bacterial expression cloning approach (Fig. 1A). We isolated RNA from active tardigrades as well as those that had entered the desiccation-tolerant tun state. RNA was reverse transcribed to cDNA and cloned into a bacterial expression vector. The pool of expression plasmids was transformed into *E. coli* BL21 AI. Protein expression was induced in populations of bacteria, and then bacteria expressing tardigrade proteins were then subjected to desiccation – with generally less than one in a million cells surviving – or maintained as a control group. After rehydration, recovery, and overnight outgrowth of the surviving bacteria, plasmids were isolated, and the cDNA inserts were amplified by linear PCR. We then sequenced the resulting DNA to determine enrichment of tardigrade genes in control and desiccated samples. 43.9 to 51.7% of annotated *Hypsibius exemplaris* transcripts and 51.0 to 58.5% of annotated *Ramazzottius varieornatus* transcripts were detected in each control library (Fig. S1A,B), suggesting we screened approximately 50% of predicted transcripts for each species. For libraries derived from the tardigrade *H. exemplaris*, 118 cDNAs were enriched in desiccated samples and 111 genes were enriched in controls (Fig. 1B, Fig S1D). The cDNAs enriched in desiccation comprise a substantial portion of the pool that was sequenced, indicative of selection (Fig. 1C,E). For libraries derived from the tardigrade *R. varieornatus*, 385 cDNAs were enriched following desiccation, and 284 were enriched in controls (Fig. 1D,E, Fig. S1E).

**Figure 1.**
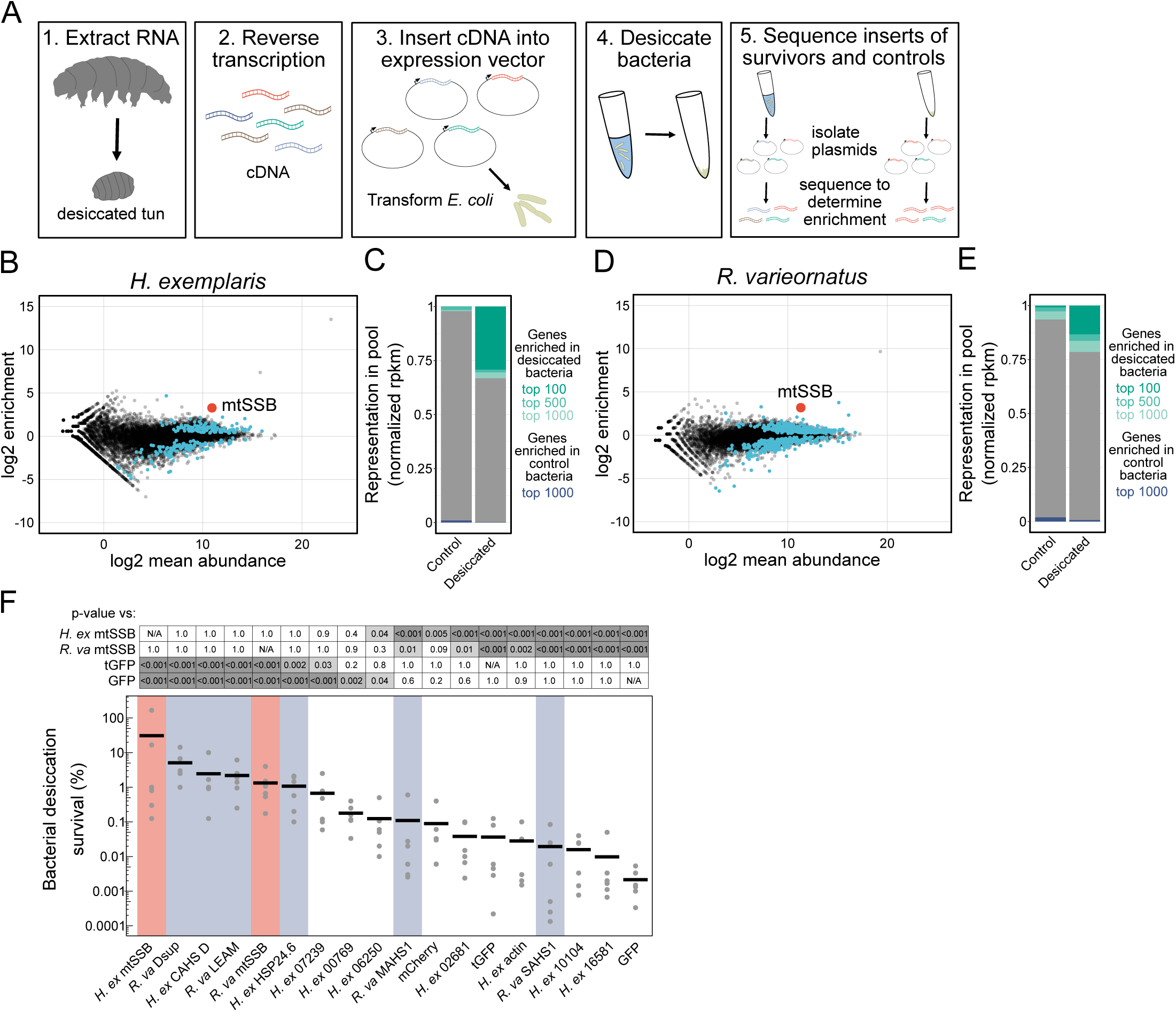
Bacterial expression cloning identified mtSSBs from *R. varieornatus* and *H. exemplaris* as potent desicco-protectants. **A)** Schematic of experimental workflow for expression cloning screens. **B)** Fold enrichment after desiccation of *H. exemplaris* genes expressed in bacteria is shown vs. log2 mean abundance (cpm). mtSSB is highlighted. **D)** Fold enrichment after desiccation of *R. varieornatus* genes expressed in bacteria is shown vs. log2 mean abundance (cpm). mtSSB is highlighted. Other genes that were significantly enriched in control or desiccated samples in B and D are indicated with blue points. **C,E)** Stacked bar plots show representation of the most enriched genes in each library in desiccated and control samples. The top 100, 500, and 1000 genes that were enriched in desiccated samples are indicated along with the top 1000 genes that were enriched in control samples. These graphs depict the extent of selection after desiccation. **F)** Desiccation survival of bacteria expressing mtSSBs (red), known protectants (blue), and negative control proteins. There was a significant difference across conditions (p<0.001, 1-way ANOVA, n=6). P-values of some pairwise comparisons from a Tukey test are shown (n=6).

We found that a cDNA encoding a mitochondrial single-stranded DNA-binding protein (mtSSB) was among the most enriched genes following desiccation for the library derived from *H. exemplaris*, and that a cDNA encoding an mtSSB was among the most enriched genes from *R. varieornatus* as well (Fig. 1B,D). To determine the extent to which mtSSBs are bona fide desicco-protectants, we compared their effectiveness in bacteria with some of the most effective tardigrade protectants reported in the literature to date, including CAHS D, Dsup, a mitochondrial late embryogenesis abundant (LEA) protein, and a small heat shock protein, HSP24.6^12–16^. We also included a number of putative control cDNAs, including transcripts that were downregulated in dried tardigrades (Fig. S2A), actin, and fluorescent proteins, which further served as a visual control for protein expression (Fig. S2B). Expression of each tardigrade mtSSB improved desiccation survival over 100-fold compared to several control proteins, and to levels that were statistically indistinguishable from the strongest known tardigrade desicco-protectants (Fig. 1F). Although there was some variability in expression levels and solubility of proteins (Fig. S2C) and differential effects on bacterial growth (Fig. S2D), these factors do not explain the efficacy of mtSSBs: there was no correlation between these variables and desiccation survival (Fig S2E,F). Each of the proteins expressed was codon optimized for *E. coli*, although codon optimization did not significantly impact mtSSB function (Fig. S2G).

One possible explanation for the efficacy of heterologously expressing mtSSBs is that they could activate the bacterial SOS response, an endogenous stress response pathway. In fact, overexpression of *E. coli* SSB and human SSBP1 has been shown to induce modest amounts of DNA damage^17^. We measured filamentous growth, a hallmark of the bacterial SOS response, to determine if this might explain the desiccation-protective effect^18^. The *H. exemplaris* mtSSB caused an increase in bacterial length, but the full-length version of the *R. vareiornatus* protein did not induce filamentous growth in bacteria, suggesting that its survival benefit is not primarily due to activation of the SOS response (Fig. S2H). While mtSSBs could significantly improve bacterial desiccation survival, they had no effect on survival of heat stress (Fig. S3A). However, the expression of *R. varieornatus* mtSSB did convey a modest benefit in protecting cells against ionizing radiation (Fig. S3B). Thus, mtSSBs do not universally improve stress tolerance, but demonstrate effects contingent upon the specific stress. We conclude that mtSSBs are potent desicco-protectants and may provide some cross tolerance to other DNA-damage inducing stresses.

### mtSSBs promote desiccation tolerance by binding to DNA

To determine the mechanism by which tardigrade mtSSBs promote desiccation tolerance, we focused our attention on *R. varieornatus* mtSSB because it likely does not induce the SOS response (Fig S2H). The predicted tetrameric structure of *R. varieornatus* mtSSB is similar to that of other mtSSBs as well as prokaryotic SSBs, with core oligonucleotide/oligosaccharide binding (OB) folds (Fig. 2A)^19–21^. Notably, both *H. exemplaris* and *R. varieornatus* mtSSBs lack a disordered linker region and acidic C-terminal tip that are characteristic of the endogenous *E. coli* SSB^22^. Instead, tardigrade mtSSBs have a longer disordered N-terminal tail. Because the acidic tip of the *E. coli* SSB is important for recruiting binding partners^23^, we did not think it likely that heterologous expression of mtSSBs would improve desiccation survival by recruiting endogenous bacterial DNA repair machinery.

**Figure 2.**
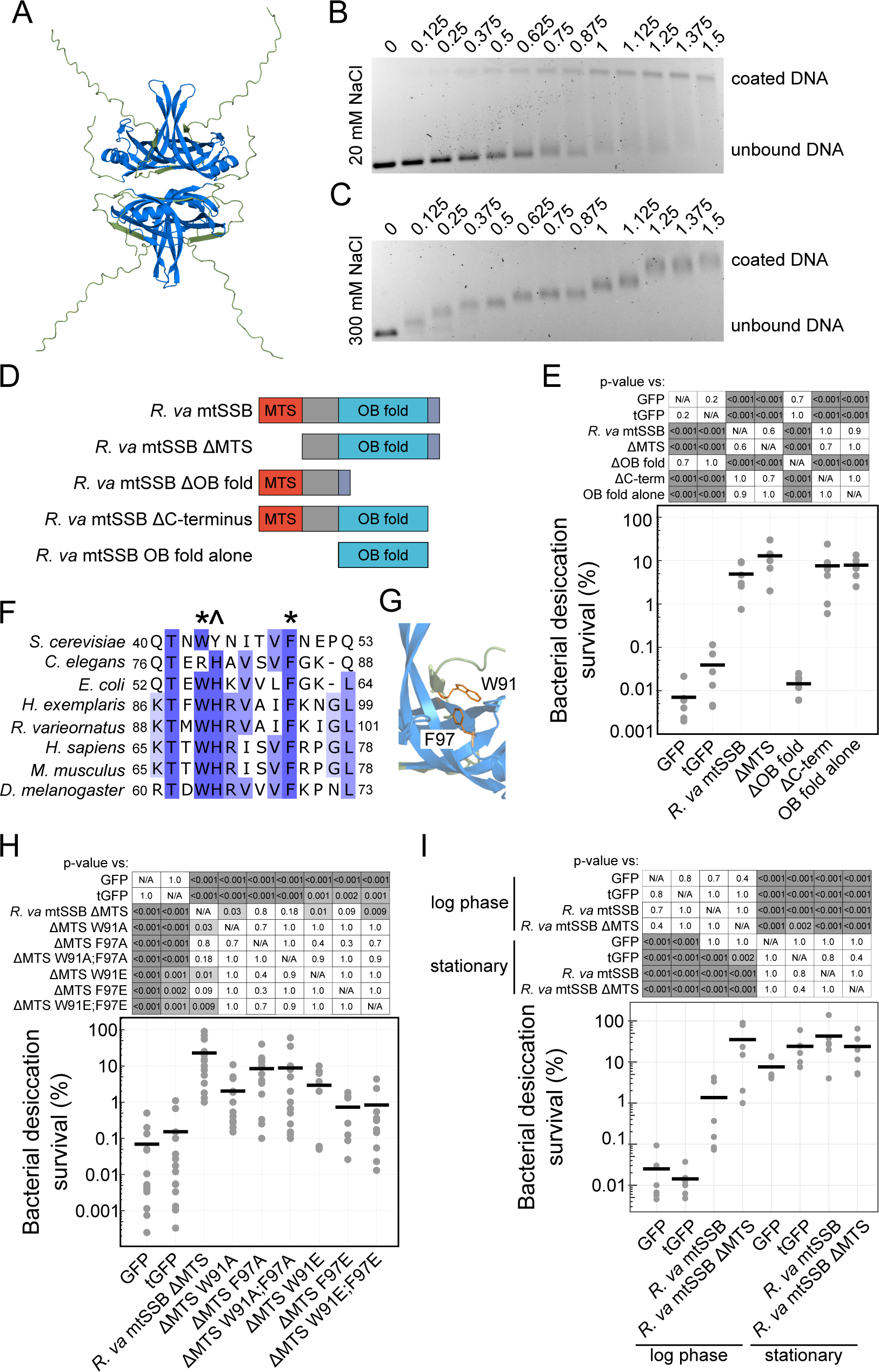
mtSSBs protect against desiccation by binding DNA. **A)** The predicted tetrameric structure of *R. varieornatus* mtSSB ΔMTS is shown. The highly conserved oligonucleotide/oligosaccharide binding (OB) fold is colored blue. **B)** Gel shift analysis of *R. varieornatus* mtSSB ΔMTS at low salt concentration (20 mM NaCl) reveals cooperative binding to ssDNA (M13mp18 circular DNA). **C)** Gel shifts conducted in buffer with 300 mM NaCl reveal incremental, non-cooperative, binding of the protein to ssDNA. **D)** A schematic of domain deletion constructs of *R. varieornatus* mtSSB. **E)** Desiccation survival of bacteria expressing *R. varieornatus* mtSSB constructs. The OB fold is necessary and sufficient for bacterial desiccation survival (n=6). **F)** A portion of sequence alignment of mtSSBs with conserved tryptophan (W) and phenylalanine (F) residues indicated with asterisks. These residues are likely important for the DNA-binding ability of the protein. The conserved histidine (H), annotated with ^, is important for dimerization of the protein. **G)** The tryptophan and phenylalanine side chains extend into the DNA-binding pocket of the OB fold and are important for stacking interactions when binding nucleotides. **H)** Desiccation survival of bacteria expressing *R. varieornatus* mtSSB ΔMTS with point mutations that convert the aromatic W or F residues to either alanine (A) or glutamic acid (E) (n=7-14). **I)** Bacteria in stationary phase were significantly more tolerant of desiccation than those in log phase. The desicco-protective effects of mtSSB expression were specific to replicating cells (n=6). P-values in E, H, and I were calculated with Tukey tests following significant 1-way ANOVA results.

However, the predicted structural conservation of *R. varieornatus* mtSSB suggested its ability to bind DNA. To directly test for DNA binding we purified *R. varieornatus* mtSSB without the predicted mitochondrial targeting sequence (ΔMTS, see Fig. S4 for MTS predictions). We assessed DNA binding by gel-shift assays, combining protein and M13mp18 ssDNA at defined ratios^24, 25^. Purified *R. varieornatus* mtSSB ΔMTS demonstrated cooperative DNA binding at a low salt concentration (20 mM NaCl), as indicated by bimodal bands in the gel (Fig. 2B). At a higher salt concentration (300 mM NaCl), binding to DNA was non-cooperative, instead showing incremental increases in the location of bands with increasing protein concentrations (Fig. 2C). These salt-dependent binding modes of *R. varieornatus* mtSSB mimic those of the *E. coli* SSB – evidence for the general conservation of the DNA-binding function of these proteins and ability to engage in both cooperative and non-cooperative interactions with ssDNA^24–26^. Non-cooperative binding is likely the predominant mode *in vivo*, especially as osmolytes become concentrated as drying cells lose water^3^.

To determine if DNA binding was essential for the desicco-protective function of mtSSBs, we tested the ability of constructs containing different regions of the protein to protect desiccated cells (Fig. 2D-E). The mitochondrial targeting sequence and C-terminal region were dispensable for protection (Fig. 2E, Fig. S5A). The OB fold was necessary and sufficient to strongly improve survival of desiccated bacteria (Fig. 2E), revealing that DNA binding is likely the salient functional property for improving desiccation survival. We then leveraged previous work that has been carried out on the mechanisms of SSB-DNA binding to test this hypothesis more specifically. Protein sequence alignments revealed conservation of the tryptophan (W) and phenylalanine (F) that have been shown to play an important role in DNA binding (Fig. 2F, Fig. S6)^27^. These aromatic amino acids extend into the ssDNA-binding pocket of the OB fold and stabilize ssDNA by binding with base-stacking interactions (Fig. 2G)^19^. Mutations in these residues reduced the protective ability of the proteins, with a substitution to negatively charged glutamic acids (E) of either site generally providing a more significant effect than substitution with likely less disruptive alanines (A) (Fig. 2H, Fig. S5B). Together, these data support the conclusion that DNA binding is essential for the desicco-protective ability of mtSSB.

Single-stranded DNA is exposed by normal cellular functions like replication and transcription, as well as during genotoxic stress^28^. To determine if the protective effects of mtSSB were cell-cycle dependent, we assessed the protective capacity of mtSSB in log and stationary phase cells. Cells in stationary phase were generally more tolerant of desiccation, as has been observed in bacteria and yeast^29–32^. As seen before, expression of *R. varieornatus* mtSSB (+/-MTS) in replicating cells improved desiccation survival; however, there was not a significant effect of mtSSB overexpression in stationary phase cells (Fig. 2I). Interestingly, it has been reported that genomes of stationary phase *E. coli* have more exposed ssDNA than cells in log phase^28^. Collectively, this could indicate that ssDNA exposed at replication forks may be important for these effects or that other protective mechanisms of stationary phase cells may diminish the added benefit of mtSSB expression.

### Tardigrade mtSSBs can localize to mitochondrial DNA and promote yeast desiccation survival

In bacteria, tardigrade mitochondrial SSBs may function similarly to endogenous SSBs, with access to genomic DNA. However, in eukaryotic cells mtSSBs localize to mitochondria, and the replication protein A (RPA) complex functions analogously in the nucleus. The tardigrade mtSSBs we identified have predicted mitochondrial targeting sequences (Fig. S4). To determine if tardigrade mtSSBs can localize to mitochondria, we added GFP tags for visualization and expressed the proteins in *S. cerevisiae* (Fig. 3A). The full-length versions of both *R. varieornatus* and *H. exemplaris* mtSSBs exhibited punctate localization that overlapped with an endogenous mitochondrial Cox4::mCherry tag (Fig. 3B). Expression of the mitochondrial targeting sequence (MTS) from each protein fused to GFP resulted in more diffuse localization throughout mitochondria. Deleting the MTS from each protein resulted in cytosolic localization, without discernable punctae. We confirmed these localization patterns in yeast stained with Mitotracker (Fig. S7). We hypothesized that the punctate distribution of mtSSBs within the mitochondria could represent binding to mitochondrial DNA (mtDNA). Indeed, in cells stained with DAPI, mtSSB punctae overlapped with DAPI-labeled mtDNA (Fig. 3C,D). Thus, tardigrade mtSSBs expressed in yeast likely associate with mtDNA.

**Figure 3.**
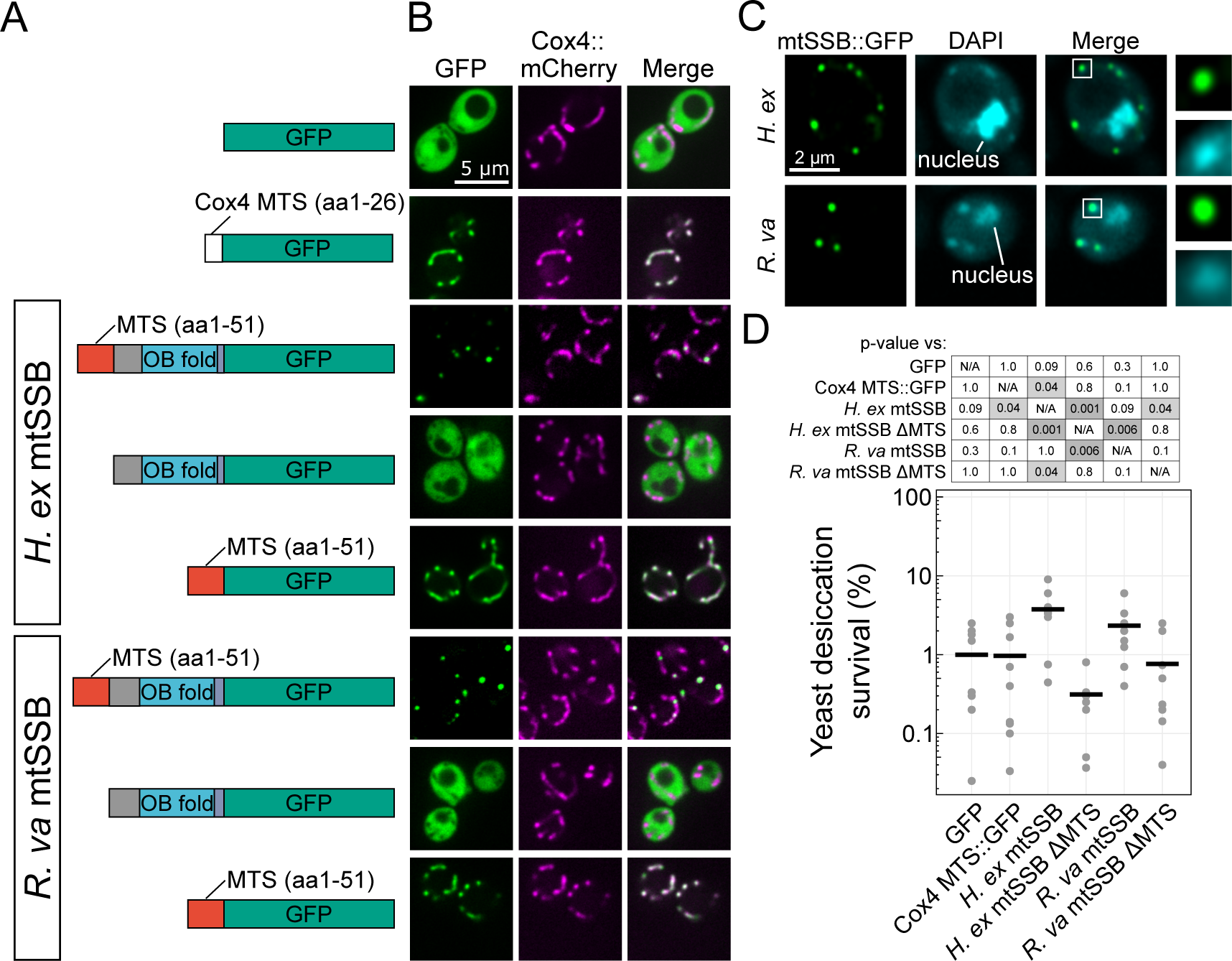
mtSSBs can enter mitochondria and protect yeast from desiccation. **A)** Schematic of yeast expression constructs used to assess protein localization. **B)** Co-expression of GFP-tagged proteins with endogenous Cox4::mCherry shows mitochondrial localization of tardigrade mtSSBs. **C)** *H. exemplaris* and *R. varieornatus* mtSSBs co-localize with DAPI-stained mtDNA. Boxes within merged images indicate regions shown to the right at 3.08x magnification. **D)** Mitochondrial localization is required for mtSSBs to improve yeast desiccation survival. P-values reported were calculated with a Tukey test (n=9).

While mtSSBs were highly effective in protecting desiccating bacteria, it was unclear if they would demonstrate the same potency in a eukaryotic cell with its diverse set of organelles, when localized only to mitochondria. We compared desiccation survival of yeast expressing tardigrade mtSSBs in different compartments to determine if localization of mtSSBs in mitochondria could also serve a protective role. Yeast expressing *H. exemplaris* mtSSB localized to mitochondria had slightly improved desiccation survival compared to yeast expressing the cytosol-localized version of the same protein (Fig. 3D). This suggests that this mtSSB can confer at least some protection when localized to mitochondria, and *in vivo*, mtSSB might function as a protectant in mitochondria in concert with other protectants functioning in other organelles.

### The ability to protect cells from desiccation is a conserved property of the ssDNA-binding domain of some SSBs

Our finding that tardigrade mtSSBs’ single-stranded DNA-binding activity makes them potent desicco-protectants suggested to us that mtSSBs from other organisms would share this potency. mtSSBs are well-conserved proteins (Fig. 4A). Eukaryotes have both mitochondrial and nuclear SSBs: typically, one copy of an mtSSB and one copy each of three nuclear replication protein A (RPA) subunits (RPA1, 70kDa; RPA2, 32 kDa; and RPA3, 14kDa). One exception is *C. elegans*, which does not have a homolog of the RPA3 subunit, but instead has another RPA2 paralog, RPA-4^33^. Expression of mtSSB proteins (either full length or ΔMTS) from tardigrades (*R. varieornatus* and *H. exemplaris*), fruit flies (*D. melanogaster*), nematodes (*C. elegans*), mouse (*M. musculus*), and yeast (*S. cerevisiae*) was indeed sufficient to increase bacterial desiccation survival (Fig. 4B, Fig. S8). Tardigrade mtSSBs were among the most potent of those tested. Human mtSSB1 was only sufficient to improve desiccation tolerance when lacking its MTS (Fig. 4B, Fig. S8C). In general, full length and ΔMTS proteins affected bacterial desiccation survival similarly, although some differences, like with *C. elegans* MTSS-1 (full length vs. ΔMTS, p<0.001, Tukey test) and *H. exemplaris* mtSSB (full length vs. ΔMTS, p=0.01, Tukey test), could reflect differences in protein folding, stability, or DNA-binding affinity in a cleaved vs. uncleaved state^34^. Although the MTS sequences are predicted to be cleaved, we do not yet know if this happens *in vivo* in tardigrades. The OB fold in isolation from many mtSSBs could also improve desiccation survival (Fig. S8C,D).

**Figure 4.**
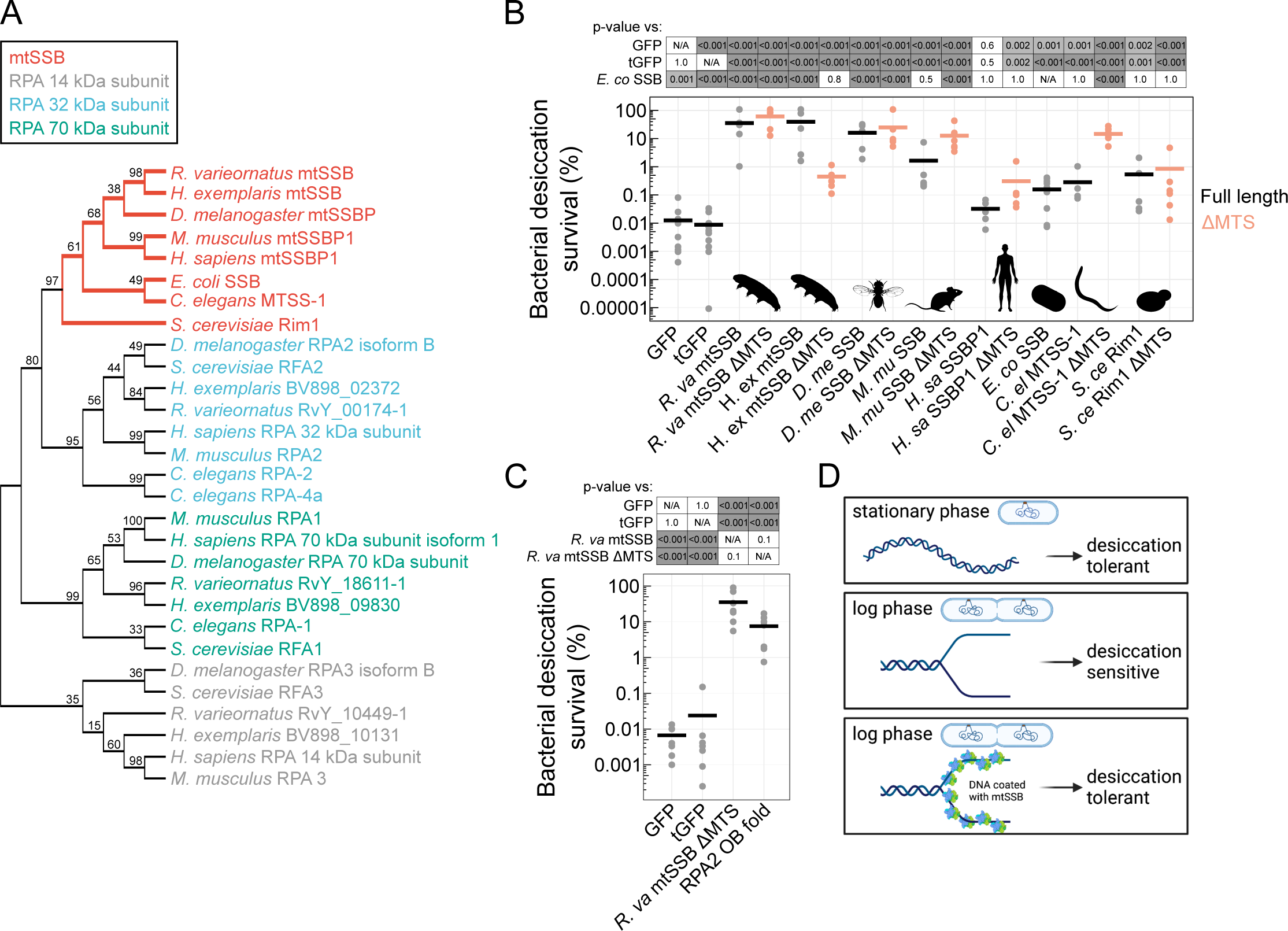
Coating DNA with protein is a conserved mechanism for desicco-protection. **A)** A maximum likelihood gene tree depicts relationships between protein sequences of homologs of replication protein A subunits and mtSSBs from multiple species. Numbers at branch points indicate bootstrap values. **B)** Desiccation survival of bacteria expressing mtSSBs from different species with and without mitochondrial targeting sequences (n=5-11). **C)** Expression of the consensus OB fold from RPA2 (NCBI ref) was sufficient to improve bacterial desiccation tolerance (n=7). P-values in B and C were derived from Tukey tests. **D)** A summary model for the role of mtSSBs in promoting desiccation tolerance.

*S. cerevisiae* Rim1 has weaker dimerization affinity than other mtSSBs due to the presence of a tyrosine instead of a highly conserved histidine at the dimer interface (Fig. 2F)^35, 36^. This provided a specific hypothesis that the reduced efficacy of Rim1 to promote bacterial desiccation tolerance relative to *R. varieornatus* mtSSB may be due to reduced dimerization affinity (Fig. 4B, p<0.001). Indeed, a version of the *R. varieornatus* mtSSB carrying a H92Y mutation had reduced desiccation survival relative to the WT protein (Fig. S9A,B). Other amino acid substitutions to replace H92 of *R. varieornatus* mtSSB produced similar effects (Fig. S9C,D). However, reverting the Y44 of *S. cerevisiae* Rim1 to H was not sufficient to significantly improve desiccation survival, suggesting that other sequence changes also contribute to differences in the potency of these proteins as desicco-protectants (Fig. S9A). Nonetheless, dimerization affinity likely impacts the level of desicco-protection conferred by mtSSBs, perhaps through altered DNA binding affinity or other properties dependent on tetramer assembly^35^.

Interestingly, overexpression of the endogenous *E. coli* SSB had a significantly weaker effect than many of the proteins from other organisms (Fig. 4B), despite strong expression and solubility (Fig. S8A,B). It is possible that the effect of *E. coli* SSB is already saturated by endogenous expression of the protein, or that overexpression could lead to dysregulation of SSB binding partners that contribute to essential processes like replication and the DNA-damage response. The non-native proteins and the OB folds of these proteins alone lack the binding sites to recruit *E. coli* SSB binding partners. Because these proteins still protected against the lethality of desiccation, we conclude that DNA-binding of these proteins improves desiccation tolerance without recruitment of functional binding partners.

We next inquired if eukaryotic nuclear SSBs (RPAs) could promote desiccation tolerance. RPAs assemble as heterotrimers to carry out similar functions to mtSSBs. They also contain OB fold domains that facilitate DNA binding. None of the RPA subunits from either *H. exemplaris* or *R. varieornatus* was significantly enriched in our screen, perhaps due to limited potency of individual subunits without assembly of the heterotrimeric complex or the fact that mitochondrial proteins like mtSSB may already be poised to function in bacteria. To determine if the nuclear analog of mtSSBs harbor similar potency as desiccation protectants, we measured desiccation survival of bacteria expressing a consensus OB fold from the RPA2 family (consensus protein sequence from cd03524, NCBI). The generalized RPA2 OB fold was sufficient to improve desiccation survival to a level that was indistinguishable from *R. varieornatus* mtSSB ΔMTS (Fig. 4C, Fig. S8E). This offers further support for the model that generic coating of ssDNA can contribute to protecting desiccating cells.

## Discussion

We set out to screen for novel protectants from tardigrades in a way that biased towards protectants that could function in heterologous contexts and would not rely on criteria like endogenous transcriptional upregulation, shared biochemical properties with known protectants, or sequence similarity to known protectants. Our bacterial expression cloning screens identified tardigrade mtSSBs as conserved proteins that dramatically improved bacterial desiccation survival. Indeed, this approach revealed a function for mtSSBs as desicco-protectants even though they are not transcriptionally upregulated during desiccation (Fig. S10)^14, 37^. As such, a strength of our expression cloning approach is its effectiveness while remaining agnostic to the functions of proteins in the source organism. In this way, expression cloning screens like the one we developed present an efficient method to identify functional properties of proteins even from emerging model organisms that are not amenable to traditional forward genetic screening. It also allows the discovery of potent protectants that don’t rely on their native environments and binding partners to function. While mtSSBs were strong desicco-protectants in bacteria, in yeast this effect was muted (Fig. 3D). The increased complexity of eukaryotic cells, with many subcellular compartments, will likely limit the potency of some protectants that work well in prokaryotes. Expression cloning screens carried out in eukaryotic cells may reveal more protectants and perhaps some that can work in the context of different subcellular compartments to protect cells from extreme stress.

We demonstrate that DNA binding is essential for mtSSBs to confer desicco-protection (Fig. 2). SSBs are non-specific in binding ssDNA^26^. Therefore, we envision a mechanism similar to what has been proposed for Dsup: protein can coat DNA to limit damage during genotoxic stresses, including desiccation^13, 38–40^. The potency of Dsup has been demonstrated in a number of contexts, yet it has also been reported to alter normal transcriptional regulation and may itself promote DNA damage in some contexts^40–42^. This suggests that the presence of DNA-coating proteins like Dsup or mtSSBs may be advantageous during extreme stress but may prove disruptive under normal physiological conditions. mtSSBs differ from Dsup in their affinity for single-stranded DNA versus a preference for nucleosome-bound dsDNA^38^. While ssDNA is generally prone to damage^43^, the efficacy of mtSSBs raises the question of whether ssDNA may be particularly vulnerable during desiccation in bacteria and perhaps more generally. ssDNA breaks are a prominent form of DNA damage in tardigrades during short term desiccation, while dsDNA breaks occur over longer time scales^44, 45^. Our results build a model that mtSSBs (or any fragment thereof that can sufficiently bind DNA) improve desiccation tolerance by binding ssDNA (Fig. 4D). We found an effect that is specific to replicating cells, possibly due to the presence of ssDNA exposed at active replication forks. DNA wrapped around mtSSBs could be physically buffered against DNA damage and ensuing lethality. However, we cannot rule out other properties of SSBs, like phase separation, or effects on RNA, that could also contribute to desiccation tolerance^46–48^. It is conceivable that such a mechanism could take place in either mitochondria or nuclei of eukaryotic cells to promote desiccation tolerance. Further understanding mechanisms by which single-stranded DNA-binding activity can act as a potent contributor to extreme desiccation tolerance will be of interest.

## Methods

### Expression library construction

To create bacterial expression libraries, we first extracted RNA from *Hypsibius exemplaris* (Z151) and *Ramazzottius varieornatus* (YOKOZUNA-1). For *H. exemplaris*, both active tardigrades and some that were conditioned at 97.5% relative humidity to induce tun formation were combined before isolating RNA. For *R. varieornatus*, which does not exhibit a significant transcriptional response to desiccation^37^, RNA was isolated only from active animals. RNA was extracted using a PicoPure RNA isolation kit (Applied Biosystems, Cat #KIT0204). A Cloneminer II cDNA Library Construction Kit (Invitrogen, Cat #A11180) was used to generate cDNA and clone it into plasmids for bacterial expression according to the specifications in the manual. Briefly, the first strand of cDNA was synthesized from RNA by reverse transcription (SuperScript III RT), the second strand was synthesized with *E. coli* DNA Polymerase I, and T4 DNA Polymerase created blunt-ended cDNA. attB adapters were ligated onto cDNA with T4 DNA ligase. Adapter-ligated cDNAs were fractionated on 1 mL Sephacryl S-500 HR resin columns to remove residual non-ligated adapters. Adapter-ligated cDNAs were then cloned into pDONR222 with BP clonase II. The entry libraries contained an estimated 4.6 million clones (*H. exemplaris*) and 2.7 million clones (*R. varieornatus*).

The donor vectors carrying tardigrade cDNA inserts were transformed into ElectroMAX DH10B T1 Phage Resistant Cells using an Electroporator 2510 (eppendorf) set to a voltage of 2.2 kV. After 1 hr of recovery, transformants were frozen at -80 °C. To isolate donor plasmids, cells were thawed and grown up in 50 mL LB with Kanamycin selection (50 µg/mL). Plasmids were harvested with a HiPure Plasmid Filter Midiprep Kit (Invitrogen, Cat #K210014). The library transfer reaction with LR Clonase II was carried out to move inserts into pDEST17 (Invitrogen, Cat #11803012) to generate an expression library. The expression libraries contained an estimated 1.7 million clones (*H. exemplaris*) and 25.7 million clones (*R. varieornatus*).

### Screening expression libraries

Expression libraries were transformed into *E. coli* BL21 AI (Invitrogen, Cat #C607003). 150 ng of plasmid DNA was transformed into a total of 300 µL of competent cells for expression screens. Two independent library transformations were conducted. In each case, after initial recovery for 30 min in SOC, transformed pools of bacteria were grown overnight in LB with ampicillin. Cultures were diluted 1:20 into LB with ampicillin and 0.2% L-arabinose to induce protein expression and split into different cultures at this stage for technical replicates. After 4 hr of induction, for each technical replicate 3 mL of culture was collected for a control sample and 3 mL of culture was collected for desiccation. Plasmids were isolated from controls with a Purelink HQ Mini Plasmid Purification Kit (Invitrogen, Cat #K2100-01). Bacteria that were to be desiccated were concentrated by centrifugation (5,000 rpm, 10 min), washed with 0.85% NaCl, and retained as a pellet. Tubes were desiccated overnight in a Savant speedvac concentrator and then kept for four days in a desiccation chamber with drierite desiccant. After desiccation, bacteria were rehydrated and allowed overnight outgrowth of survivors in LB with ampicillin. As with controls, plasmids were isolated from these populations of desiccation survivors with minipreps (Invitrogen, Cat #K2100-01).

### Sequencing

To capture biological and technical variability in this screen, we sequenced two technical replicates (i.e., two controls and the paired desiccated samples) from each of the two independent library transformations. To avoid sequencing primarily pDEST17 plasmid backbone, the cDNA inserts of the plasmids were isolated and amplified by low-cycle (12x) PCR using a Kapa2G Robust Hotstart PCR kit (Roche, Ref #07961073001) according to specifications. Primer sequences were CATCACCATCACCTCGAATCAAC and TTCGGGCTTTGTTAGCAGCCTCGAATC. The annealing temperature for the reaction was 55 °C and the extension time was 2 minutes. The protocol for the kit suggests 15 s/kb, thus allowing for amplification of inserts up to ∼8 kb. These PCR amplicons were cleaned with a ChargeSwitch-Pro PCR cleanup kit (Invitrogen, Cat #CS32050). Amplicons were sonicated to ∼200 bp and sequencing libraries were prepared by the UNC high-throughput sequencing facility (HTSF) using the Kapa DNA hyper kit. Batches were pooled onto lanes for paired end 75 bp read sequencing on a HiSeq4000.

### Sequence analysis

Quality of reads from sequencing was assessed with fastqc. Paired reads were mapped to the genomes of *H. exemplaris* (v3.1.5) or *R. varieornatus* (Rv1) with bowtie2^37^. Mapped reads were assigned to genes with featurecounts. Genes without any reads mapped to them were removed prior to determining enrichment in control or desiccated conditions with DESeq2^49^.

Genes with an FDR <0.05 were considered to be significantly enriched in desiccated or control samples. The integrative genomic viewer (IGV) was used to visualize read alignment to portions of the genome^50^.

### Identification of tardigrade mtSSBs

The identity of Rv07050 and BV898_11351 as mtSSBs was determined by reciprocal BLAST. The *H. exemplaris* genome (v3.1.5) has another annotated transcript, BV898_04979, with 94.6% sequence identity to the *H. exemplaris* mtSSB, BV898_11351 (Fig. S1C)^37^.

However, BV898_04979 had very limited coverage of library reads mapping to it, and we were unable to amplify it from either cDNA or genomic template (Fig. S11,S12). Furthermore, expression of the hypothetical BV898_04979 protein did not confer the same desiccation survival benefits as the tardigrade mtSSBs (Fig. S13), suggesting that the annotated sequence differences of BV898_04979 reduce its function, at least in this heterologous context. Thus, we believe that *H. exemplaris*, like most other species, has a single mtSSB. Because BV898_04979 and BV898_11351 are near the ends of genomic scaffolds, it is possible that these scaffolds represent a contiguous genomic region with a single mtSSB.

### Molecular cloning

To clone mtSSBs and other control proteins, sequences were codon optimized for *E. coli* (Integrated DNA Technologies). To clone non codon-optimized versions of *R. varieornatus* mtSSB and *H. exemplaris* mtSSB, RNA was isolated from tardigrades, converted to cDNA using the SuperScript III First-Strand Synthesis System (Invitrogen, Cat #18080-051) with oligoDT priming, and PCR amplified from total cDNA with gene-specific primers including 30bp of homology to pDEST17. pDEST17 was linearized by high fidelity PCR (Q5 High-Fidelity 2x Master Mix, NEB, Cat #M0492), during which sequence encoding the 6x His tag was removed. Sequences of primers used to linearize pDEST17 were TGATTCGAGGCTGCTAACAAAG and GTAGTACGACATATGTATATCTC. PCR products were gel-extracted and cleaned (Zymo Gel DNA Recovery Kit, Cat #D4002), and assembled into the pDEST17 backbone with NEBuilder HiFi Assembly Master Mix (NEB, Cat #E2621). Correct assembly of each plasmid was verified by sequencing (Genewiz/Azenta).

Other versions of proteins like mtSSB domain constructs and species’ ΔMTS versions were similarly assembled. To clone plasmids for expression of the OB fold domain of mtSSBs from various species, the NCBI conserved domain search tool was used to identify the OB fold of each protein^51^. The region of each protein with homology to cd04496 (SSB_OBF) was cloned into pDEST17 for bacterial expression. To create expression constructs with individual amino acid substitutions, plasmids were edited via site-directed mutagenesis (Q5 Site-Directed Mutagenesis Kit, NEB, Cat #E0554).

### Bacterial expression of proteins

Heterologous expression of proteins was carried out in *E. coli* BL21 AI (Invitrogen, Cat #C607003). Single colonies were grown overnight at 37 °C in LB with ampicillin selection.

Overnight cultures were diluted 1:20 into LB with ampicillin and 0.2% L-arabinose (Thermo scientific, Cat #A11921.18) to induce protein expression. Expression of GFP and mCherry was visually apparent, providing a simple visual confirmation of protein induction. To assess protein expression levels, protein was visualized with SDS-PAGE and Coomassie staining. Bacteria that were induced for 4 hr at 37 °C with 0.2% L-arabinose were pelleted in 1.5 mL tubes by centrifugation at 5,000 rpm for 10 minutes and resuspended in 200 µL of 0.85% NaCl. Bacteria were lysed with 30 pulses of sonication using a Branson Sonifier 250 set to 50% output and 50% duty cycle. To isolate the soluble fraction, total lysate was centrifuged at 14,000 rpm for 10 minutes at 4 °C. Protein concentration in each fraction was determined with a Bio-Rad Protein Assay Kit (#5000002). 2 µg protein was combined with sample loading buffer (2x buffer = 4 % SDS, 20% glycerol, 0.125 M Tris-HCl pH 6.8, 10% 2-mercaptoethanol, 0.004% bromphenol blue), heated at 95 °C for 10 minutes, and loaded into wells of a NuPAGE 4-12% Bis-Tris mini protein gel (NP0322BOX). Samples were run alongside protein marker (Precision Plus Protein Kaleidoscope, Bio-Rad #1610375). Samples were run in MOPS SDS Running Buffer (Invitrogen, NP0001) for 70 minutes at 140 V. Gels were stained in Coomassie (0.1% Coomassie brilliant blue, 50% methanol, 40% water, 10% acetic acid) overnight at 4 °C with rocking and destained in a 5:4:1 solution of water:methanol:acetic acid. Gels were imaged with a Bio-Rad Universal Hood II Molecular Imager using Image Lab 5.2.1 software. The expected molecular weights of proteins that are reported were computed with the Expasy Compute pI/Mw tool^52^. Quantification of relative protein expression or solubility (Fig. S2E) was carried out in FIJI by subtracting the background of gel images and taking the ratio of the integrated density of the band for each protein divided by the total integrated density of the lane.

### Analysis of bacterial growth rate and length

We determined bacterial growth rates by measuring OD600 over time in a plate reader. Overnight cultures of bacteria were diluted 1:20 for induction in LB with ampicillin and 0.2% L-arabinose in a 96-well plate. Plates were kept at 37 °C in a BioTek Synergy H1 plate reader with double orbital shaking. OD600 measurements were taken every 10 min. Three independent biological replicates were conducted. Length of bacteria was determined by imaging cells on a Nikon Eclipse E800 microscope with a pco.panda sCMOS camera and measuring lengths in FIJI.

### Bacterial desiccation, heat, and radiation survival

The OD600 of bacteria was measured after 4 hrs of induction with L-arabinose. The volume of bacteria to be desiccated was normalized to a concentration of 1.5 OD600 for each condition. Bacteria were collected by centrifugation (5,000 rpm, 10 min), washed with 1 mL 0.85% NaCl, and resuspended in 1 mL 0.85% NaCl. A dilution series was plated onto LB+AMP plates to determine the concentration of bacteria in controls. Bacteria were pelleted by centrifugation (5,000 rpm, 10 min) and the supernatant was removed before overnight desiccation in a Savant speedvac concentrator. Bacteria were rehydrated in the same volume, and a dilution series was plated to determine the concentration of survivors. Survival was calculated as the cfu after desiccation divided by that of controls. To assess survival of heat shock and radiation, the same protocol was followed except with exposure to 52 °C for 1 hr or to 2,180 Gy irradiation.

### Yeast expression and imaging

Expression constructs carrying mtSSBs, or parts thereof, were codon optimized for yeast. These fragments were synthesized (IDT) with overhangs for BP reactions (Invitrogen BP clonase II, Cat #11789-020) to insert constructs into pDONR221. Subsequent LR reactions (Invitrogen LR clonase II, Cat #11791-020) moved constructs from pDONR221 into pAG415. Cloning reactions were carried out per kit instructions. Plasmids were sequence-verified to ensure correct assemblies.

Plasmids were transformed into *S. cerevisiae* (W303) MATa ura3-1, leu2-3,112, his3-11, trp1-1, can1-100, ade2-1. Briefly, a 5 mL overnight culture of yeast was grown in yeast bacto-peptone dextrose (YPD) at 24 °C in an orbital shaker. 0.5 mL of overnight culture was added to 25 or 50 mL of culture and grown to log phase (OD660 0.3-0.8) with shaking. Yeast cells were pelleted by centrifugation and resuspended in 5 mL of TE Lithium acetate. Cells were again concentrated by centrifugation and most of the supernatant was removed. Yeast in minimal remaining supernatant were kept on ice for 15 min. Salmon sperm DNA (Agilent technologies, Cat #201190) was boiled for 5 min and then stored on ice. For each transformation, 50 µL of cells, 5 µL of salmon sperm DNA, and 1 µL of expression plasmid were combined in an Eppendorf tube. To each tube, 250 µL of TE/lithium acetate/PEG was added. Samples were incubated at 32 °C with shaking for 30 min and heated to 42 °C for 15 min. Cells were then pelleted by centrifugation, resuspended in 250 µL 1M sorbitol, and plated onto SD -LEU plates for selection.

The endogenous Cox4::mCherry reporter strain was created by fusing endogenous Cox4 sequence to an mCherry::kanMX selectable marker (pFA6) by PCR. The following primers were used to amplify this sequence: GAATGTGGTTCTGTTTACAAACTAAACCCTGTTGGTGTTCCAAATGATGACCACCATCACcggatccccgggttaattaa,AATAAAGAAGAAGGTAAAAAGTAAAAGAGAAACAGAAGGGCAACTTGAATGATAAGATTAGaattcgagctcgtttaaac. The PCR product was purified (Zymoclean Gel DNA Recovery Kit, Cat #D4002) and transformed into *S. cerevisiae* W303 following the same protocol as with plasmids, except that cells were kept on a roller drum at 24 °C for 18 hrs outgrowth in YPD before plating on YPD with 2x G148 for selection.

To visualize protein localization, yeast were grown overnight in SG -LEU with adenine (0.5 mg/mL) for induction (24 °C with shaking). Yeast not carrying the Cox4::mCherry reporter were stained with Mitotracker Red FM (Invitrogen, M22425) to confirm localization patterns. Cells induced overnight in SG -LEU with adenine were isolated and stained in 50 µM Mitotracker Red and 0.1 M HEPES in yeast extract plus casamino acids (YC complete, no sugar) for 30 min. Cells were washed and resuspended in YC complete without sugar for imaging. Cells with GFP- and mCherry-labeled proteins or Mitotracker staining were imaged with a Nikon TiE base with CSU-X1 spinning disk head (Yokogawa), and ImagEM EMCCD camera (Hamamatsu) using Metamorph software for image acquisition.

To stain yeast with DAPI, cells were first washed in YC complete without sugar. Cells were then fixed in 4% paraformaldehyde (Electron Microscopy Sciences, Cat #15710) in PBS for 1 hr with rocking. Cells were mounted on glass slides in a 1:1 ratio of YC complete:DAPI Fluoromount-G (SouthernBiotech, Cat #0100-20). Yeast were imaged with a Zeiss 880 LSM with fast Airyscan.

### Yeast desiccation survival

To measure desiccation survival, tardigrade mtSSBs without fluorescent labels and control proteins were transformed into yeast. Protein expression was induced and mid-log phase (OD660 0.4-0.6) cells were concentrated for desiccation. Dilution series’ of controls and samples after overnight desiccation in a speedvac and 1 hr rehydration in SD -LEU liquid media were plated (onto SD -LEU) to determine cfu concentrations. Survival was calculated as the ratio of cfu after desiccation divided by that of controls.

### Sequence alignment and MTS prediction

Predictions of mitochondrial targeting sequences from mtSSBs were generated with TargetP2.0^53^. Protein sequences for mtSSBs and *E. coli* SSB were aligned with MUSCLE. The predicted mitochondrial targeting sequences were removed for alignments. The alignment was visualized with Jalview and colored according to sequence identity.

### Protein structure prediction

The structure of *R. varieornatus* mtSSB ΔMTS was predicted using alphafold colab v2.3.2^21^. The multimer model was used with an input of four copies of the *R. varieornatus* mtSSB ΔMTS. We also confirmed that introduction of DNA-binding mutations to W91 and F97 did not significantly disrupt predicted protein structure.

### Protein purification

Sequence encoding 6X His::TEV was added to the bacterial expression construct carrying *R. va* mtSSB ΔMTS by PCR amplification and ligation (Q5 Site-Directed Mutagenesis Kit, NEB, Cat #E0554). The plasmid insert was sequence-verified and then transformed into *E. coli* BL21 AI (Invitrogen, C607003). Single colonies were picked into 50 mL LB with ampicillin for overnight culture at 37 °C with shaking. Overnight cultures were diluted 1:20 into a total volume of 2 L of LB with ampicillin and 0.2% L-arabinose. Protein expression was induced for 6 hr, at which point bacteria were harvested by centrifugation (5,000 rpm, 10 min, at 4 °C). Cell pellets were washed in 0.85% NaCl and frozen at -80 °C.

The cell pellet was thawed in Talon buffer A (50 mM sodium phosphate pH 7.4, 300 mM NaCl, 5 mM imidazole) with 50 µg/ml lysozyme, 12.5 U/mL DNase, and 1x protease inhibitors (Roche). Resuspended cells were sonicated then centrifuged at 12,000 xg for 30 min. Soluble lysate was 0.45 µm filtered then loaded onto 1 mL Talon Crude columns (Cytiva). After washing to baseline with buffer A, bound protein was eluted with Talon buffer B (50 mM sodium phosphate pH 7.4, 300 mM NaCl, 250 mM imidazole).

Eluate from Talon affinity chromatography was collected and dialyzed against 20 mM Tris HCl pH 7.4, 150 mM NaCl. TEV protease was added at a 10:1 target:protease mass ratio while in dialysis and left to incubate at 4 °C overnight. The digest at this stage had a volume of approximately 3 mL and was diluted to approximately 25 mL in Talon buffer A to facilitate column loading. The samples were loaded on 1 mL Talon Crude columns with the flowthroughs collected as ’released protein.’ Flowthroughs were concentrated to approximately 500 µL, and all samples were analyzed with SDS PAGE.

### Gel shift assays

Gel shift assays were carried out as previously described^24, 25^. Purified *R. varieornatus*

ΔMTS mtSSB was combined with 4.7 nM M13mp18 ssDNA (NEB, N4040S) at ratios calculated with the following equation: R = n x ([mtSSB tetramer]/[nucleotide]), using n=33 nucleotides/tetramer. Protein and DNA were incubated together for 1 hr at 22 °C in 30 μL of buffer T (10 mM Tris (pH8.1), 0.1 mM EDTA) with additions of either 20 mM NaCl or 300 mM NaCl. 2 μL of loading dye (50% glycerol, 0.04% bromophenol blue) was added to each reaction and loaded into wells of a 0.5% agarose gel. Gels were run in 20 mM Tris, 0.4 mM sodium acetate, 0.2 mM EDTA running buffer for 3 hrs at 60 V, then soaked in buffer T with 1 M NaCl for 1 hr at 4 °C. Gels were stained for 30-60 minutes in 2 μg/mL ethidium bromide and destained for 1 hr in buffer T with 1 M NaCl, also at 4 °C. Gels were imaged with a Bio-Rad Universal Hood II Molecular Imager and Image Lab 5.2.1 software. Images were inverted for display in figures (Fig. 2B,C).

### Phylogenetic tree construction

MEGAX was used to create a phylogenetic tree of replication protein A (RPA) and mtSSB homologs^54, 55^. The following protein sequences were obtained from NCBI: NP_002936.1, NP_001284487.1, NP_002938.1, XP_005250105.1, AAH19119.1, AAH04578.1, AAH28489.1, NP_001273592.1, NP_524274.1, NP_001260652.1, NP_001285131.1, NP_536744.2, KZV13472.1, CAA96241.1, CAA89468.1, NP_009958.2, WP_089642251.1. The following protein sequences were obtained from Wormbase: *C. elegans* RPA-1, RPA-2, RPA-4a, and MTSS-1. The following tardigrade homologs were identified via BLAST and protein sequences obtained from *H. exemplaris* genome annotation v3.1.5, and *R. varieornatus* genome Rv1: BV898_09830, BV898_02372, BV898_10131, BV898_11351, RvY_18611-1, RvY_00174-1, RvY_10449-1, RvY_07050-1^37^. Full length sequences (including mitochondrial targeting sequences of mtSSBs) were aligned with MUSCLE. The evolutionary history was inferred by using the Maximum Likelihood method and JTT matrix-based model^56^. The percentage of replicate trees in which the associated taxa clustered together in the bootstrap test (500 replicates) are shown next to the branches^57^. Initial tree(s) for the heuristic search were obtained automatically by applying Neighbor-Join and BioNJ algorithms to a matrix of pairwise distances estimated using the JTT model, and then selecting the topology with superior log likelihood value. There were 859 positions in the final dataset.

### Statistical analysis

Significant enrichment from expression cloning screens was determined by analysis with DESeq2^49^. For survival experiments in bacteria and yeast, we first performed a 1-way ANOVA to test for differences across all conditions. If this produced a significant result, we then conducted post-hoc Tukey tests to define pairwise differences. The number of independent biological replicates is indicated in figure legends or in the text. Statistical analyses were carried out in Rstudio.

## Acknowledgements

We thank Jim Keck, Kevin Slep, Dorothy Erie, and members of the Goldstein lab for helpful suggestions and constructive feedback. We thank Diana Cook for advice and training with yeast culture and transformations. Nate Nicely of the UNC Protein Expression and Purification core facility at UNC assisted with protein purification. We thank the UNC High Throughput Sequencing Facility for sequencing and are grateful for advice on experimental design and analysis from Tristan De Buysscher and Piotr Mieczkowski. Kazuharu Arakawa kindly shared cultures of *R. varieornatus*. Fig. 4D was created with BioRender.com. This work was supported by the NSF (IOS 2028860, awarded to BG) and the NIH (F32GM131577, awarded to JDH).

## Author Contributions

Conceptualization, JDH and BG; Methodology, JDH, CMCH; Formal Analysis, JDH; Investigation, JDH and CMCH; Resources, KSB; Writing – Original Draft, JDH; Writing – Review and Editing, JDH, CMCH, KSB, BG; Visualization, JDH; Supervision, KSB, BG; Funding Acquisition, JDH, BG.

## Notes

### Competing Interest Statement

The authors have declared no competing interest.

