## Supplementary Figures for "A bacterial expression cloning screen reveals tardigrade single-stranded DNA-binding proteins as potent desicco-protectants"

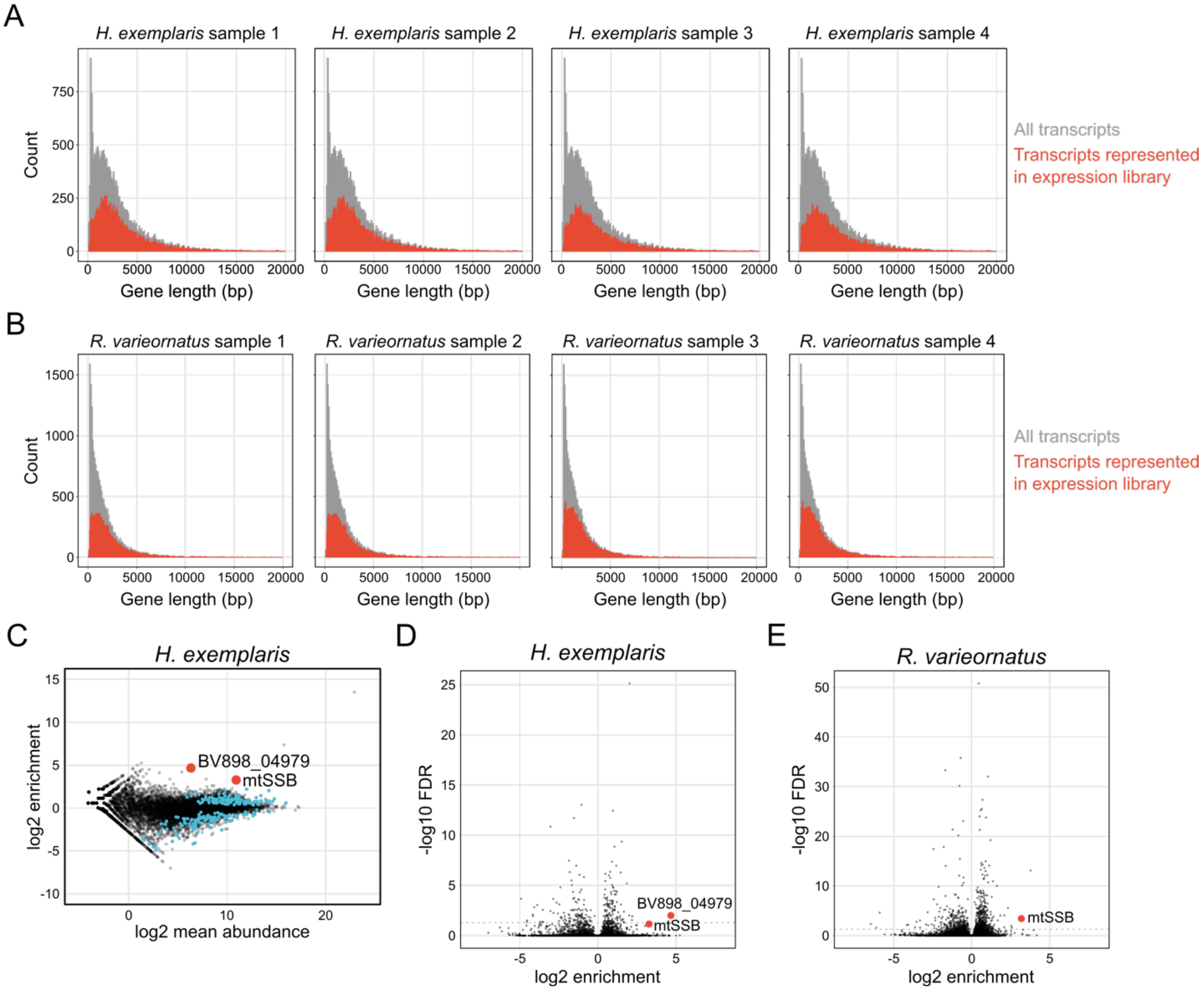


**Figure S1.** Expression cloning library representation and analysis. **A,B)** Representation of transcripts in each of four replicates of the expression libraries derived from *H. exemplaris* (A) or *R. varieornatus* (B). Any transcript with a read mapped to it is indicated in red versus the background (gray) showing all transcripts in the genome. **C)** The same plot as in Figure 1B, also highlighting BV898_04979 in addition to BV898_11351 (mtSSB). **D,E)** Volcano plots of -log10 FDR vs log2 fold enrichment of cDNAs from expression cloning screens in *H. exemplaris* (D) and *R. varieornatus* (E).

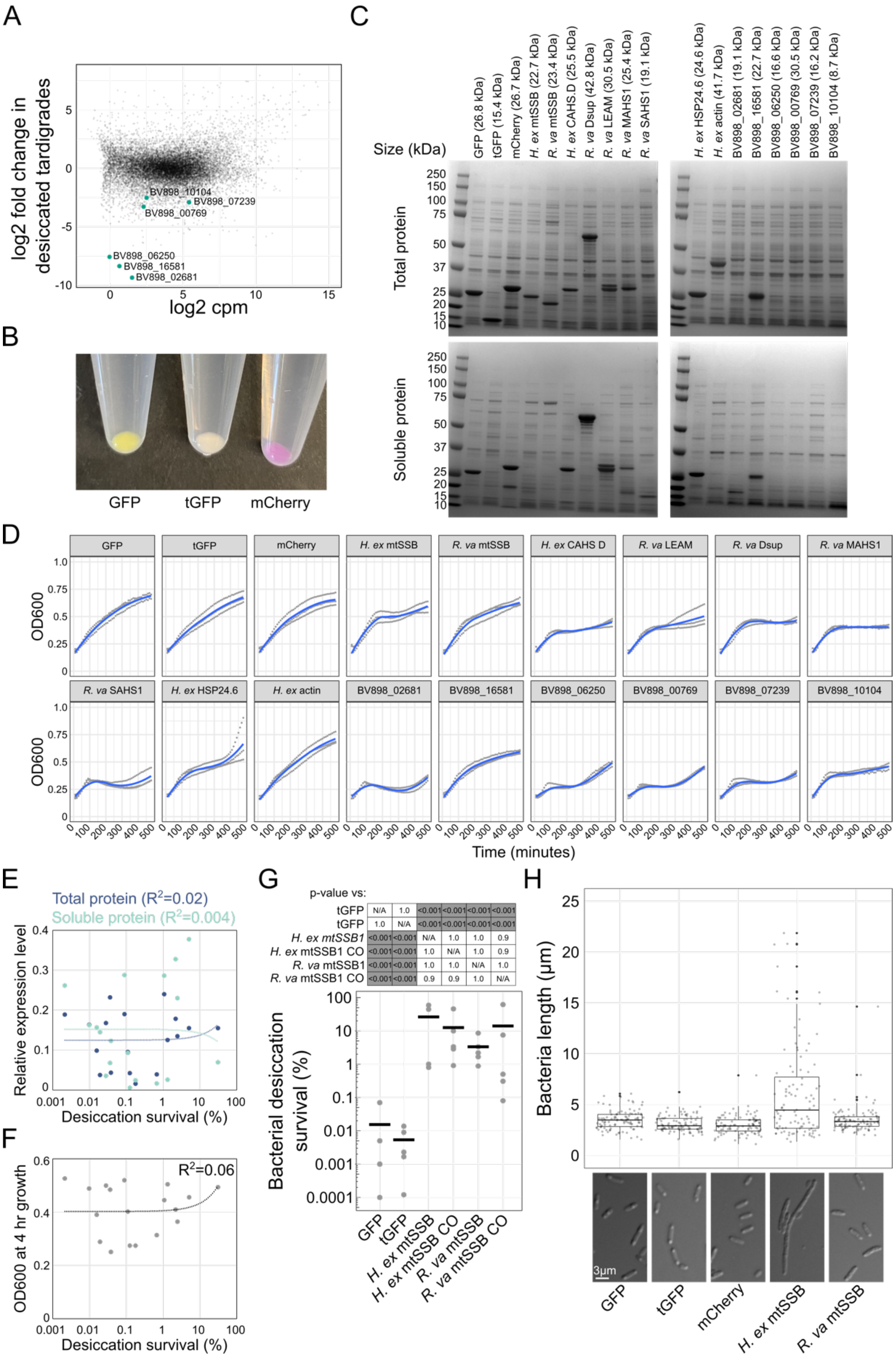


**Figure S2.** Controls for validation of tardigrade mtSSBs as desiccation protectants. **A)** A plot of transcriptional changes during desiccation in tardigrades from Boothby *et al.* 2017 was used to identify likely negative control genes that were less abundant during desiccation. **B)** Expression of fluorophores in bacteria provides a facile “tube-level phenotype” to confirm protein expression in each experiment. **C)** SDS-PAGE analysis of total and soluble protein from bacteria after 4 hrs of induction. **D)** Growth curves for 8 hrs of culture are shown for bacteria with induced expression of each heterologous protein. Three replicates are shown in gray and a fitted curve is shown in blue. Cultures were harvested at 4hr for all desiccation and control experiments. **E)** Levels of total protein expression and soluble protein in bacteria were not correlated with desiccation survival. **F)** Using OD600 after 4hr of growth as a proxy for bacterial growth, there was no correlation with desiccation survival. $R^2$ values from linear regressions are reported in E and F. Note, desiccation survival is plotted on a log axis in E and F. **G)** Comparison of codon optimized (CO) and non-codon optimized versions of *H. exemplaris* mtSSB and *R. varieornatus* mtSSB reveal negligible differences in bacterial desiccation survival. P-values reported were derived from Tukey tests. **H)** The length of bacteria heterologously expressing mtSSBs and control proteins is plotted. Filamentous growth of bacteria is a hallmark of the SOS DNA damage response. Expression of *H. exemplaris* mtSSB caused a shift towards more filamentous growth, suggesting that the SOS response may be activated.

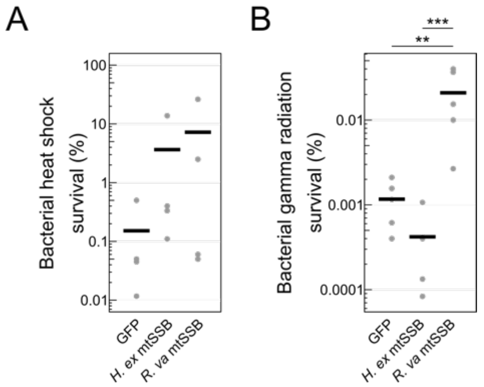


**Figure S3.** Tardigrade mtSSBs expressed in *E. coli* provide limited cross-tolerance to heat and radiation. **A)** Survival of bacteria expressing mtSSBs exposed to 52 °C heat shock for 1 hour. There were no significant differences in survival across conditions (p=0.29, 1-way ANOVA). **B)** Survival of bacteria expressing mtSSBs exposed to 2180 Gy gamma radiation. There was a significant difference in survival across conditions (p<0.001, 1-way ANOVA). Expression of the *R. varieornatus* mtSSB improved survival relative to GFP-expressing controls (p=0.002) and *H. exemplaris* mtSSB (p<0.001, Tukey test).

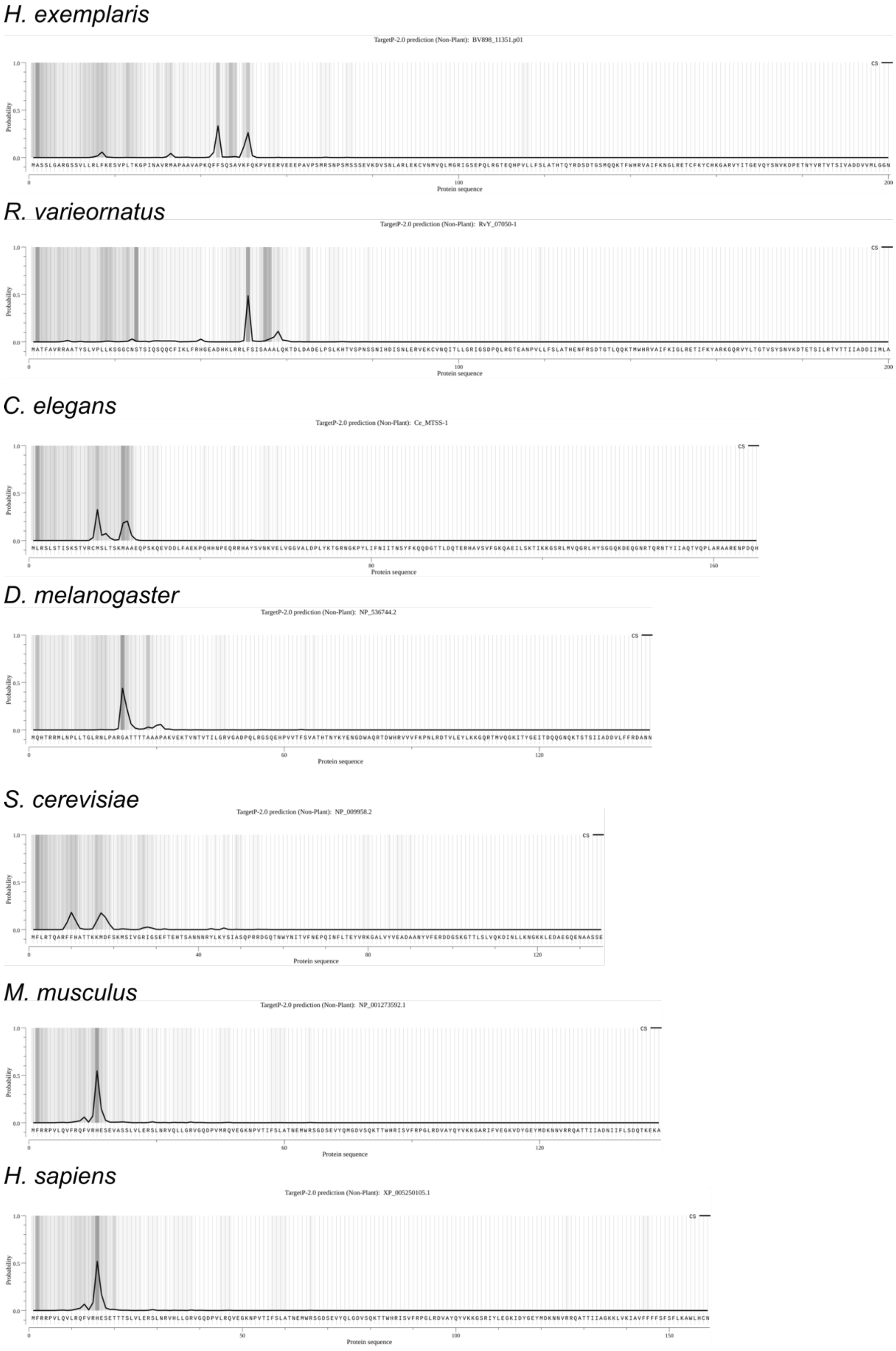


**Figure S4.** TargetP predictions of mitochondrial targeting sequences of mtSSBs from various species.

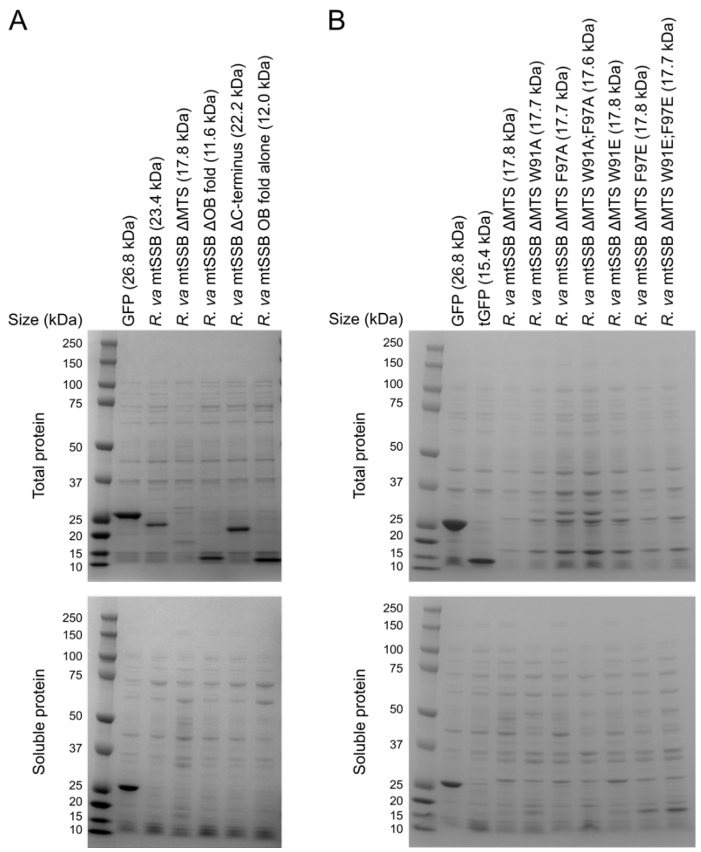


**Figure S5.** SDS-PAGE analysis of expression and solubility of heterologous expression of different mtSSB domain and mutant constructs in bacteria. **A)** Total and soluble protein from bacteria expressing domains of *R. varieornatus* mtSSB. **B)** Total and soluble protein from bacteria expressing *R. varieornatus* mtSSB with point mutations likely to disrupt DNA binding affinity.

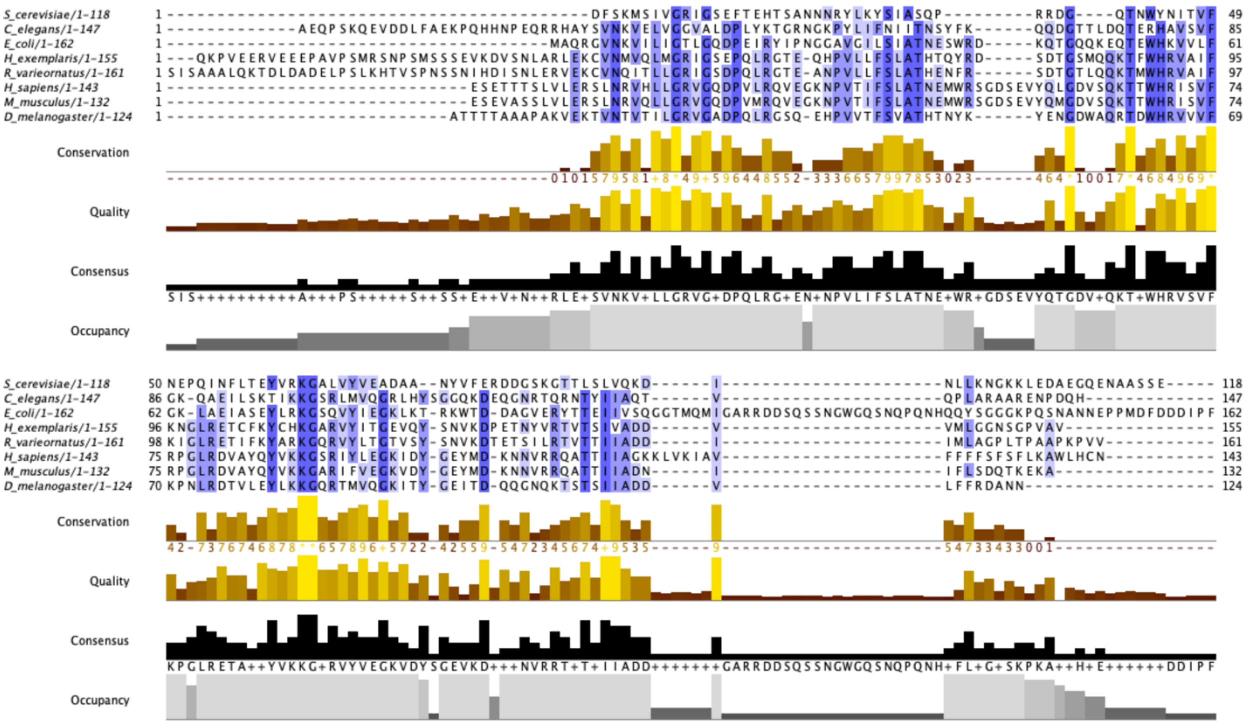

**Figure S6.** Sequence alignment of mtSSBs from various organisms as well as *E. coli* SSB. Mitochondrial targeting sequences were removed from mtSSBs. Alignment was conducted with MUSCLE and visualized with JalView.

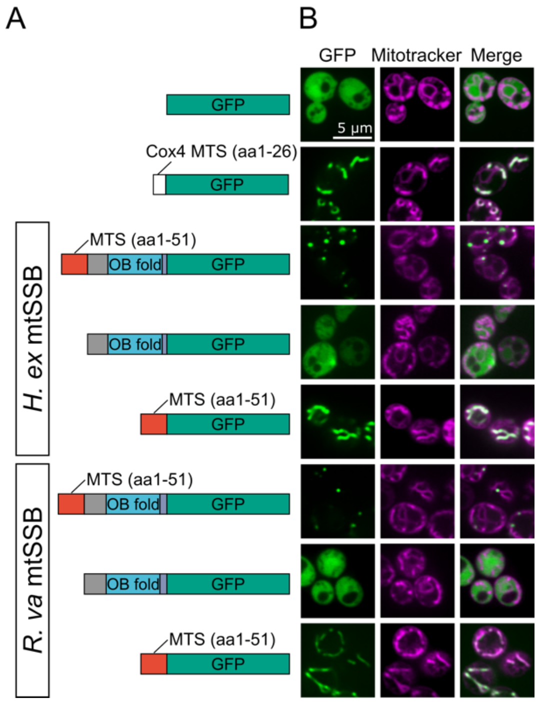


**Figure S7.** Expression patterns of fluorescent proteins and mitochondria. **A)** Diagram of expression constructs used. **B)** Expression of GFP-tagged mtSSBs and co-staining with Mitotracker reveals punctate mitochondrial localization of *H. exemplaris* mtSSB and *R. varieornatus* mtSSB.

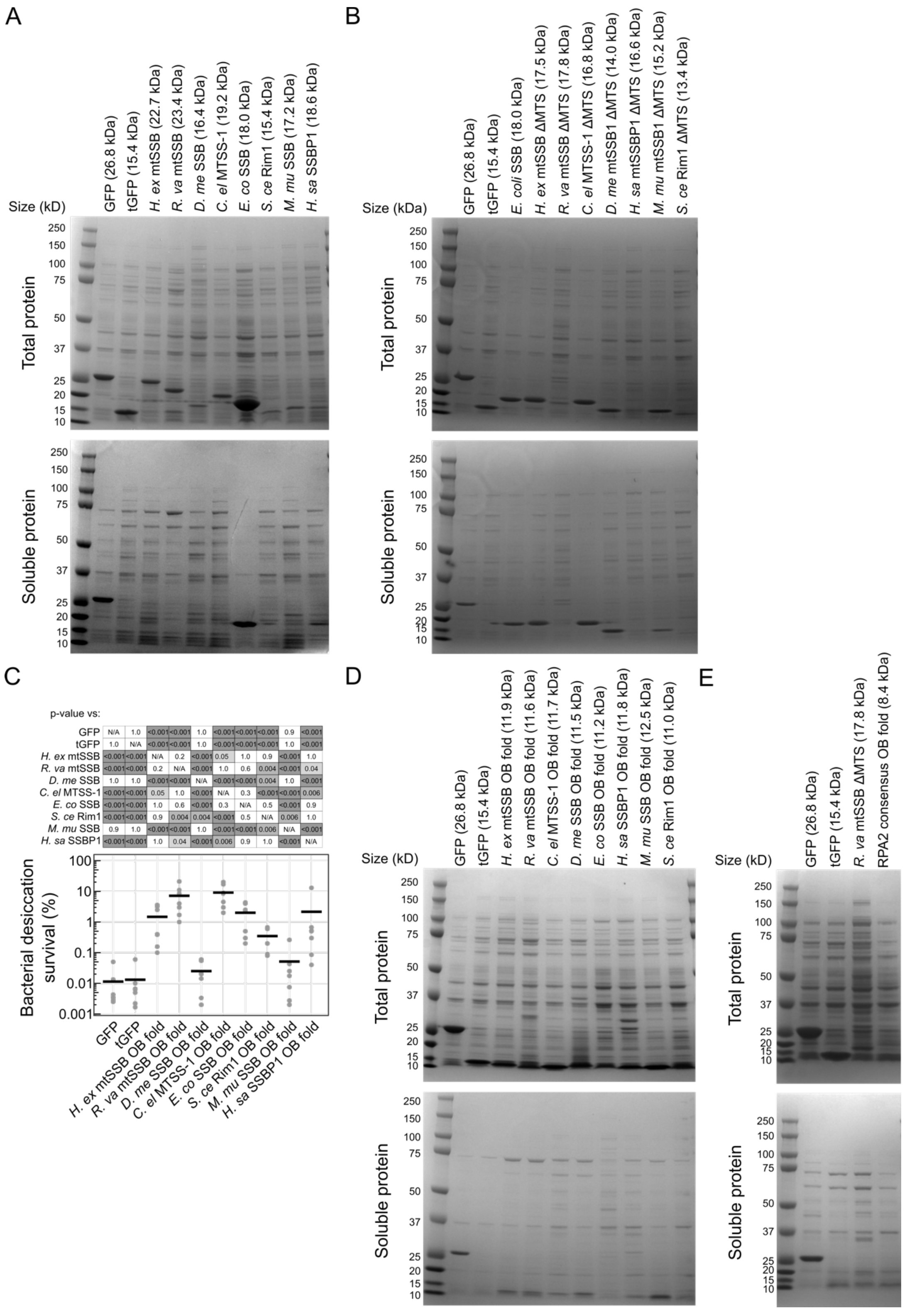


| | | | | | | | | | | |
|---|---|---|---|---|---|---|---|---|---|---|
| GFP | N/A | 1.0 | <0.001 | <0.001 | 1.0 | <0.001 | <0.001 | <0.001 | 0.9 | <0.001 |
| tGFP | 1.0 | N/A | <0.001 | <0.001 | 1.0 | <0.001 | <0.001 | <0.001 | 1.0 | <0.001 |
| *H. ex* mtSSB | <0.001 | <0.001 | N/A | 0.2 | <0.001 | 0.05 | 1.0 | 0.9 | <0.001 | 1.0 |
| *R. va* mtSSB | <0.001 | <0.001 | 0.2 | N/A | <0.001 | 1.0 | 0.6 | 0.004 | <0.001 | 0.04 |
| *D. me* SSB | 1.0 | 1.0 | <0.001 | <0.001 | N/A | <0.001 | <0.001 | 0.004 | 1.0 | <0.001 |
| *C. el* MTSS-1 | <0.001 | <0.001 | 0.05 | 1.0 | <0.001 | N/A | 0.3 | <0.001 | <0.001 | 0.006 |
| *E. co* SSB | <0.001 | <0.001 | 1.0 | 0.6 | <0.001 | 0.3 | N/A | 0.5 | <0.001 | 0.9 |
| *S. ce* Rim1 | <0.001 | <0.001 | 0.9 | 0.004 | 0.004 | <0.001 | 0.5 | N/A | 0.006 | 1.0 |
| *M. mu* SSB | 0.9 | 1.0 | <0.001 | <0.001 | 1.0 | <0.001 | <0.001 | 0.006 | N/A | <0.001 |
| *H. sa* SSBP1 | <0.001 | <0.001 | 1.0 | 0.04 | <0.001 | 0.006 | 0.9 | 1.0 | <0.001 | N/A |

**Figure S8.** Analysis of bacterial expression and solubility of mtSSBs from different species. **A)** SDS-PAGE analysis reveals expression of each full-length protein in total bacterial lysate and some variability in solubility of these proteins. **B)** SDS-PAGE analysis shows expression of mtSSBs lacking mitochondrial targeting sequences. The soluble fractions of the lysate were also run on a gel. **C)** Expression of OB fold domains from many of the proteins in Figure 4B could also improve bacterial desiccation survival, with OB fold from *R. varieornatus* mtSSB and *C. elegans* MTSS-1 being most effective. P-values were calculated with a Tukey test following a significant 1-way ANOVA ($p<0.001$). **D)** Expression of proteins from C was visualized from total and soluble fractions of bacterial lysate. **E)** Control gels for expression and solubility of the consensus RPA2 OB fold.

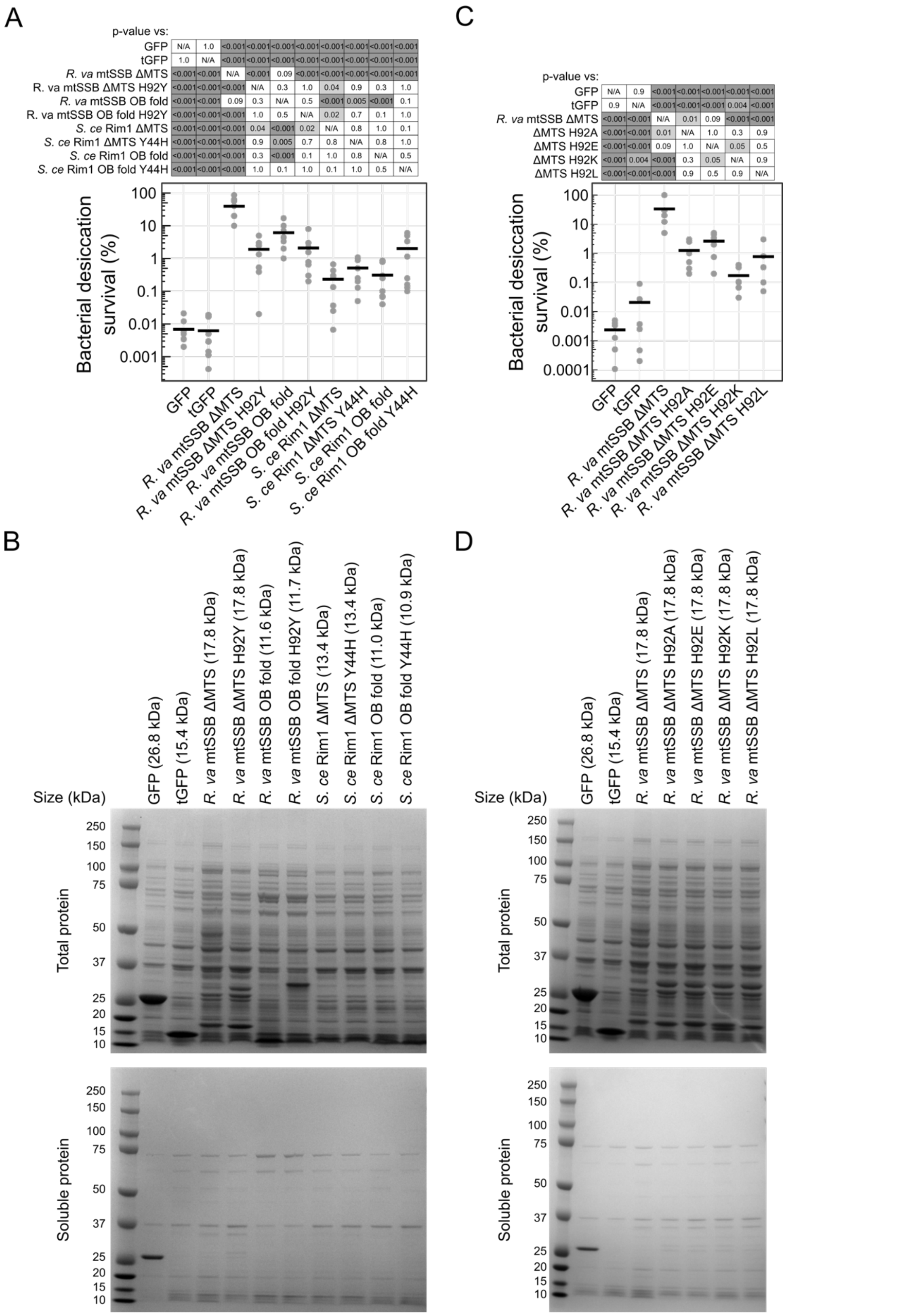


A — p-value vs:

| | | | | | | | | | | |
|---|---|---|---|---|---|---|---|---|---|---|
| GFP | N/A | 1.0 | <0.001 | <0.001 | <0.001 | <0.001 | <0.001 | <0.001 | <0.001 | <0.001 |
| tGFP | 1.0 | N/A | <0.001 | <0.001 | <0.001 | <0.001 | <0.001 | <0.001 | <0.001 | <0.001 |
| *R. va* mtSSB ΔMTS | <0.001 | <0.001 | N/A | <0.001 | 0.09 | <0.001 | <0.001 | <0.001 | <0.001 | <0.001 |
| R. va mtSSB ΔMTS H92Y | <0.001 | <0.001 | <0.001 | N/A | 0.3 | 1.0 | 0.04 | 0.9 | 0.3 | 1.0 |
| *R. va* mtSSB OB fold | <0.001 | <0.001 | 0.09 | 0.3 | N/A | 0.5 | <0.001 | 0.005 | <0.001 | 0.1 |
| R. va mtSSB OB fold H92Y | <0.001 | <0.001 | <0.001 | 1.0 | 0.5 | N/A | 0.02 | 0.7 | 0.1 | 1.0 |
| *S. ce* Rim1 ΔMTS | <0.001 | <0.001 | <0.001 | 0.04 | <0.001 | 0.02 | N/A | 0.8 | 1.0 | 0.1 |
| *S. ce* Rim1 ΔMTS Y44H | <0.001 | <0.001 | <0.001 | 0.9 | 0.005 | 0.7 | 0.8 | N/A | 0.8 | 1.0 |
| *S. ce* Rim1 OB fold | <0.001 | <0.001 | <0.001 | 0.3 | <0.001 | 0.1 | 1.0 | 0.8 | N/A | 0.5 |
| *S. ce* Rim1 OB fold Y44H | <0.001 | <0.001 | <0.001 | 1.0 | 0.1 | 1.0 | 0.1 | 1.0 | 0.5 | N/A |

C — p-value vs:

| | | | | | | | |
|---|---|---|---|---|---|---|---|
| GFP | N/A | 0.9 | <0.001 | <0.001 | <0.001 | <0.001 | <0.001 |
| tGFP | 0.9 | N/A | <0.001 | <0.001 | <0.001 | 0.004 | <0.001 |
| *R. va* mtSSB ΔMTS | <0.001 | <0.001 | N/A | 0.01 | 0.09 | <0.001 | <0.001 |
| ΔMTS H92A | <0.001 | <0.001 | 0.01 | N/A | 1.0 | 0.3 | 0.9 |
| ΔMTS H92E | <0.001 | <0.001 | 0.09 | 1.0 | N/A | 0.05 | 0.5 |
| ΔMTS H92K | <0.001 | 0.004 | <0.001 | 0.3 | 0.05 | N/A | 0.9 |
| ΔMTS H92L | <0.001 | <0.001 | <0.001 | 0.9 | 0.5 | 0.9 | N/A |

**Figure S9.** Dimerization affinity likely impacts mtSSB efficacy and is an example of a natural variation between *R. varieornatus* mtSSB and *S. cerevisiae* Rim1. **A)** Amino acid substitutions replaced histidine 92 of *R. varieornatus* mtSSB or its OB fold with the tyrosine of Rim1. Similarly, tyrosine 44 of Rim1 was swapped for a histidine. Desiccation survival of bacteria expressing these constructs is shown. **B)** Protein expression and solubility levels of proteins with amino acid substitutions were determined with SDS-PAGE. **C)** Bacterial desiccation survival is diminished in strains expressing *R. varieornatus* mtSSB with mutations in H92 that are likely to disrupt its dimerization. **D)** Total and soluble protein from bacteria expressing *R. varieornatus* mtSSB with point mutations likely to disrupt its ability to dimerize. P-values in A and C were calculated with Tukey tests.

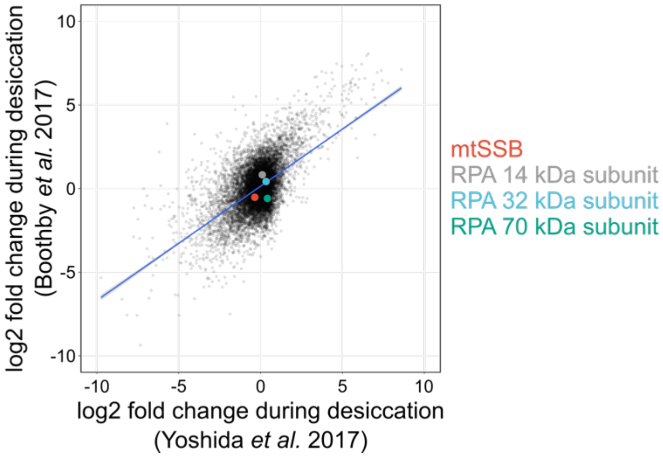


**Figure S10.** *H. exemplaris* mtSSB and replication protein A homologs do not display significant transcriptional changes during desiccation in tardigrades. Expression changes of each transcript are plotted on a graph of log2 fold change values from Boothby *et al.* 2017 and Yoshida *et al.* 2017.

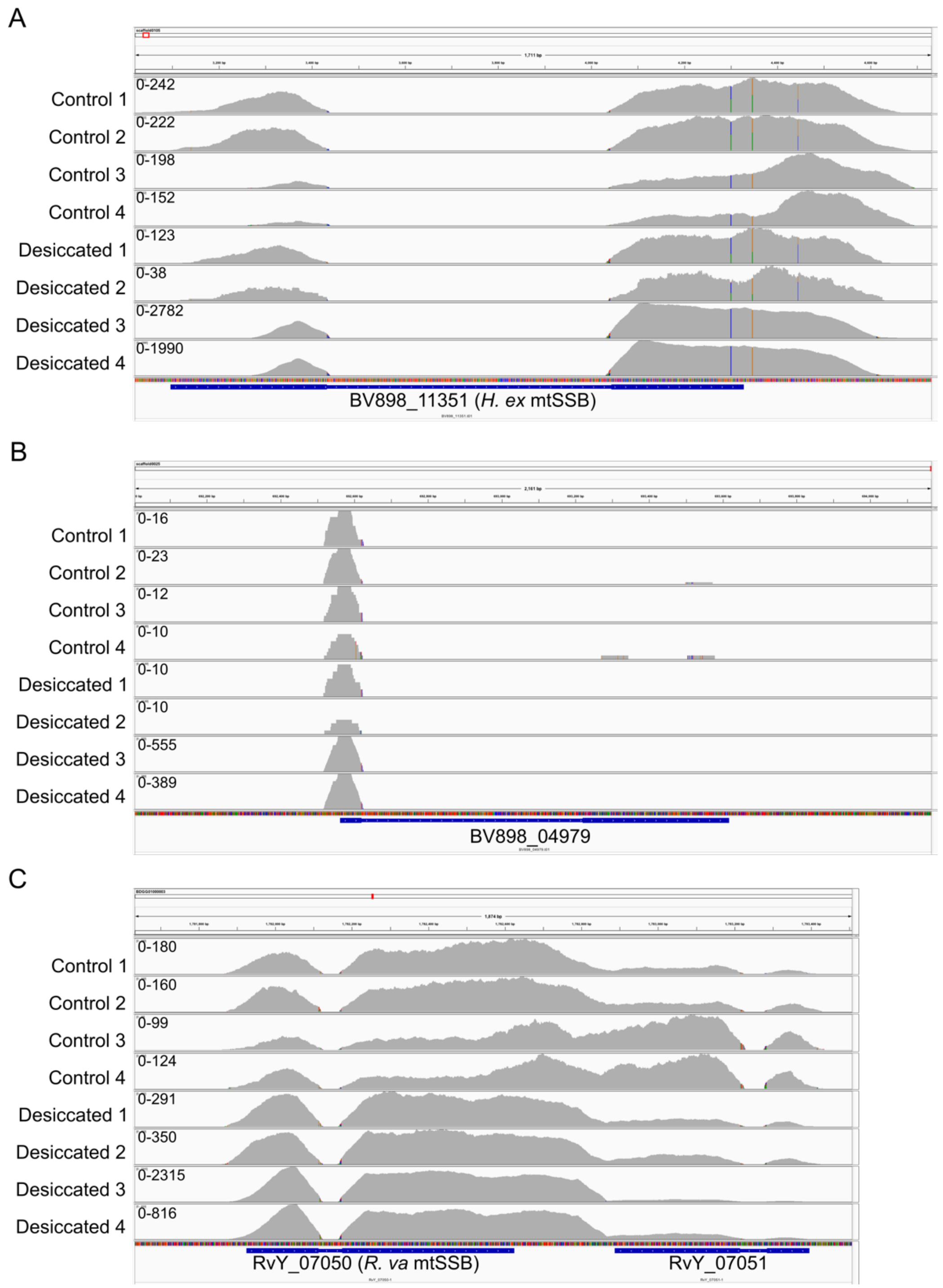


**Figure S11.** Distribution of reads aligned to BV898_04979, BV898_11351, and RvY_07050. Integrative genomics viewer was used to visualize reads aligned to each of BV898_04979 **(A)**, BV898_11351 **(B)**, and RvY_07050 **(C)**.

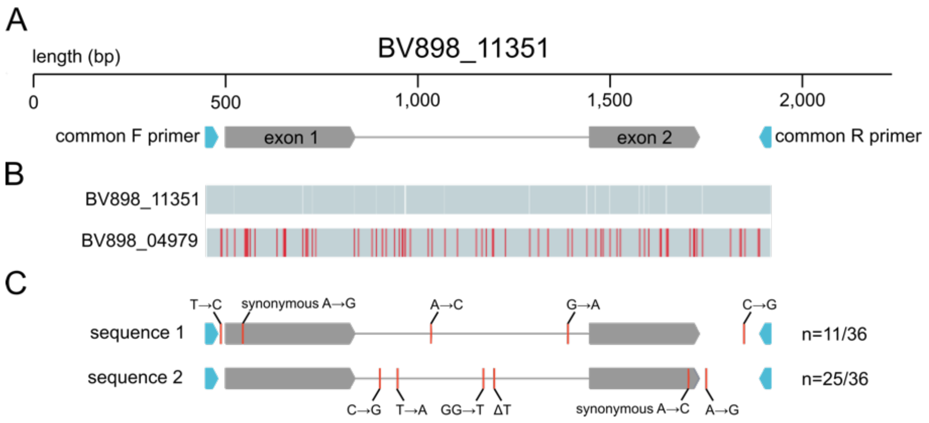


**Figure S12.** *Hypsibius exemplaris* has a single mtSSB. **A)** Annotation of the genomic locus of BV898_11351. This sequence is found on scaffold0105. BV898_11351 contains two exons. Common primers were designed in a region of identical sequences between the flanking genomic regions of BV898_04979 (scaffold0025) and BV898_11351 (scaffold0105). **B)** An alignment of genomic loci of BV898_11351 (scaffold0105) and BV898_04979 (scaffold0025) reveals significant sequence differences between the two scaffolds. **C)** Amplification and sequencing of genomic DNA from individual tardigrades revealed the presence of two versions of BV898_11351. Each of these had some polymorphisms relative to the consensus scaffold sequence. In no case was genomic sequence from scaffold0025 (BV898_04979) identified. The frequency of each sequence was derived from sequencing six clones per individual for six tardigrades.

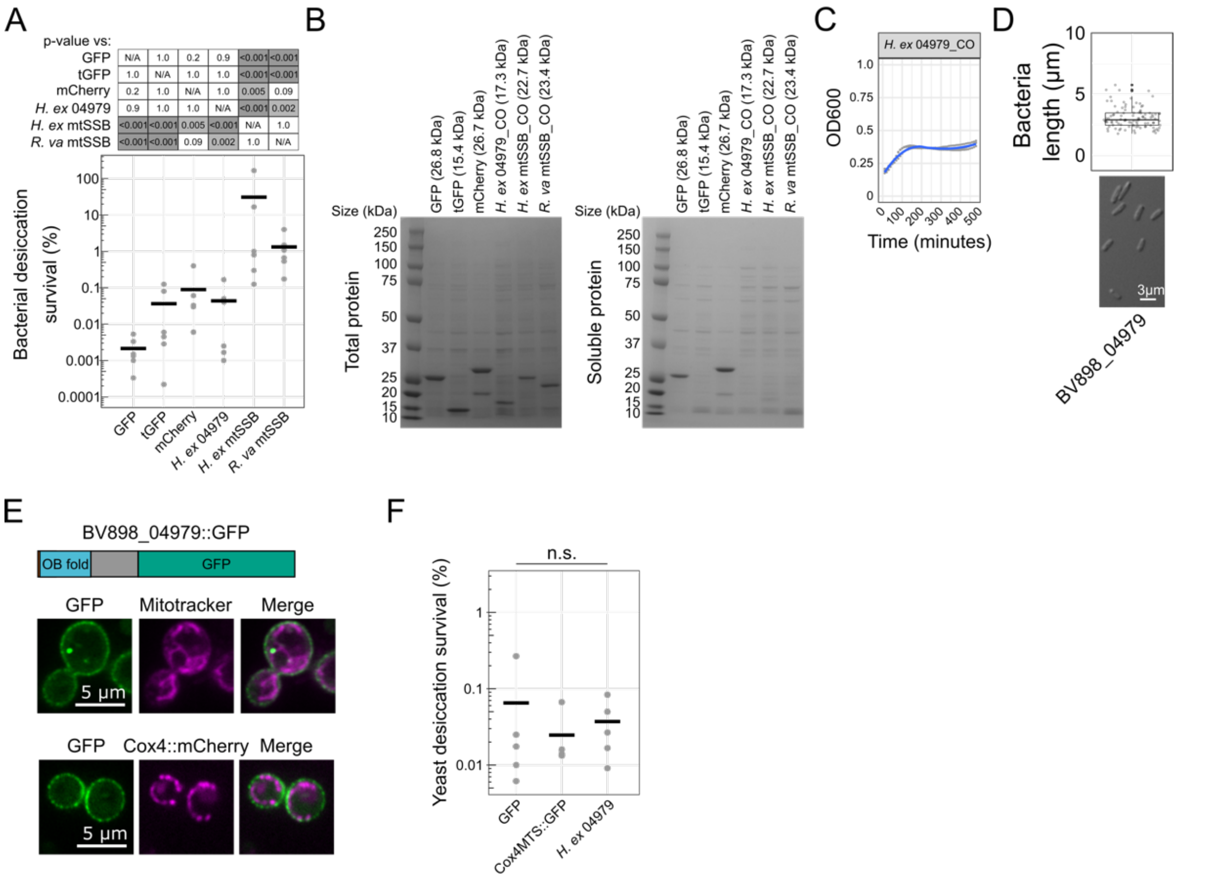


**Figure S13.** BV898_04979 serves as a negative control protein. A) Expression of BV898_04979 did not significantly improve bacterial desiccation survival. Note, data for GFP, tGFP, mCherry, *H. ex* mtSSB, and *R. va* mtSSB are the same as in Figure 1 as *H. ex* BV898_04979 was tested in parallel to these controls. The p-values reported are calculated by Tukey's test on the entire panel of strains, inclusive of *H. ex* BV898_04979. B) SDS-PAGE analysis of bacterial protein expression (total and soluble) reveals strong expression and solubility of fluorophore controls. BV898_04979 and mtSSBs were well-expressed, but had limited solubility. C) Growth analysis of *H. ex* BV898_04979-expressing bacteria reveals stunted growth relative to bacteria expressing fluorophore controls (see Figure S2D for comparison). D) Overexpression of BV898_04979 does not cause filamentous growth in BL21 *E. coli*. E) Expression of GFP-tagged BV898_04979 in yeast reveals localization towards the cell cortex. F) Expression of BV898_04979 did not significantly alter yeast desiccation survival (p=0.86, 1-way ANOVA, n=5).
